# Molecular and neuronal mechanisms governing sexually dimorphic prioritization of innate behaviors

**DOI:** 10.1101/2023.12.04.569869

**Authors:** Xinyu Jiang, Mingze Ma, Mengshi Sun, Jie Chen, Yufeng Pan

## Abstract

Males and females display dimorphic innate behaviors and further prioritize them differently. How the sexually dimorphic behavioral prioritization is mediated is poorly understood. In *Drosophila*, around 60 pairs of pC1 neurons in males and 6 pairs in females control sexually dimorphic behaviors. We show that an increase of pC1 activity determines the sequential execution of behaviors such as sex, aggression, sleep, and feeding in a sex-specific way. We identify distinct subsets of pC1 neurons in both males and females that regulate different behaviors. We further discover diuretic hormone 44 (DH44) and acetylcholine (ACh) as co-transmitters in pC1 neurons. ACh promotes the execution of each behavior in both sexes, whereas DH44 functions in a sex-specific and activity-dependent manner to establish the sexually dimorphic behavioral outputs. These findings provide a framework for understanding the molecular and cellular mechanisms underlying sexually dimorphic prioritization of innate behaviors.

## Introduction

Adapting behaviors properly, especially those innate behaviors that are born with, at any given time in a changeable world is critical for survival or reproduction but ultimately for maximum evolutionary fitness. When two or more competing stimuli pursue to access the same motor systems simultaneously, behavioral decision-making happens[1]. Such decision-making is a problem of prioritization determined by internal states, external cues, and learning from experience. Intriguingly, behavioral decision-making is often sex-specific since males and females take on distinct roles in reproduction. Males and females not only display ‘qualitatively’ (e.g., sex) and ‘quantitatively’ (e.g., sleep) dimorphic behaviors[2, 3], but also respond differently when facing challenges. For instance, pain experiences reduce the sexual motivation in female but not male mice[4]. How males and females adapt/prioritize dimorphic innate behaviors is a challenging question in any given model animal, and the hormonal and/or social modulations of these behaviors make the question even harder to explore.

*Drosophila melanogaster*, as a classic model for research on the neural basis underlying innate behaviors with great progress, offers an excellent platform for solving questions related to sexual dimorphism, given there are exclusively genetic-hardwired sex determination mechanisms without hormones unlike in mammals[5]. It is widely known that the neuronal substrates underlying sexual dimorphism in fruit flies are specified by the male-specific product encoded by *fruitless* (Fru^M^) and the sex-specific products encoded by *doublesex* (Dsx^M^ in males and Dsx^F^ in females)[6–8]. With the increasing work on diverse animals illustrating that sex differences prefer to emerge in interneurons with shared sensory and motor neurons[5, 9], the focus has been drawn to the interneurons that express *fru* and/or *dsx*. In the past decade, the significance of a subset of interneurons defined by *dsx* in the posterior brain region termed pC1 in both sexes has been recognized by more and more researchers[10–23]. There are ∼60 Dsx^M^-positive pC1 neurons in males with more than 50% co-expressing Fru^M^ (termed P1 based on *fru* expression), but only ∼6 Dsx^F^-positive counterparts in females[2, 24]. Despite the huge differences in cell numbers, they arborize and project to both shared and distinct brain regions. Nevertheless, the sexually dimorphic pC1 neurons function similarly in terms that they are thought to be a center for sexual and aggressive behaviors, integrating multiple sex-related sensory inputs and representing internal states in both males and females[24–28].

Recently, a growing body of research has revealed that P1/pC1 neurons (hereafter referred to as pC1) in male flies are involved in behavioral decision-making[13, 14, 24, 29–32]. Activating male pC1 neurons induces both courtship and inter-male aggression[11, 12]. Furthermore, the activity of pC1 neurons is, on the one hand, inhibited by sleep drive to balance sleep and sex[13, 30], and on the other hand, regulated antagonistically by sugar consumption and satiety signals [32, 33] to balance feeding and courting. Notably, evidence underlying pC1 neurons regulate courtship, sleep, and feeding in a threshold-based manner[14, 31] highlights such neurons as an arbitrating node for the behavioral prioritization in male flies. Despite great progress on neural basis underlying individual innate behaviors and their crosstalk in male flies, relatively less work has been done in females, and even less in the comparison of mechanisms about behavioral decision-making between the sexes is studied.

Here, we performed a series of experiments proving that pC1 interneurons in both sexes control a sexually dimorphic prioritization of multiple innate behaviors. We identify distinct subsets of pC1 neurons that regulate different behaviors in both sexes. Moreover, we discover the neuropeptide DH44 as a key factor in establishing the sexually dimorphic behavioral prioritization. These results reveal a neural basis underlying the flexible tuning of sexually dimorphic behavioral adaptations.

## Results

### Sexually dimorphic pC1 neurons hierarchically control multiple innate behaviors in both sexes

Previous studies have found that pC1 neurons regulate multiple innate behaviors, but these studies used different pC1 drivers and manipulations, often focusing on only one sex. To systematically explore how pC1 neurons control multiple innate behaviors in both sexes, we used an intersectional strategy as done previously (*dsx^GAL4^* and *R71G01-LexA*)[10, 34] to label homologous pC1 neurons in both sexes of the same genotype. Such intersection labeled ∼6 neurons per hemisphere in the female brain and ∼23 neurons per hemisphere in the male brain, hereafter referred to as pC1^R71G01^. We then expressed the *Drosophila* temperature-sensitive cation channel dTrpA1 in pC1^R71G01^ neurons for activation and set the temperatures as five gradients from 22℃ to 32℃, where 22℃ represented the baseline (Figure 1A). To prove that raised temperatures indeed led to a corresponding increased activity of pC1^R71G01^ neurons, we performed an *ex vivo* calcium imaging by monitoring the cell bodies of pC1^R71G01^ neurons with GCaMP6m and dTrpA1 co-expression in response to this range of temperatures. We observed that the GCaMP6m responses of pC1^R71G01^ neurons in both sexes increased as temperature rose from 22℃ to 32℃, in which males showed a roughly linear increase and females showed a slow and then fast growth trend (Figure 1B). Thus, this paradigm allowed us to compare the effects of increased activity levels of pC1^R71G01^neurons on multiple innate behaviors in both sexes.

**Figure 1.**
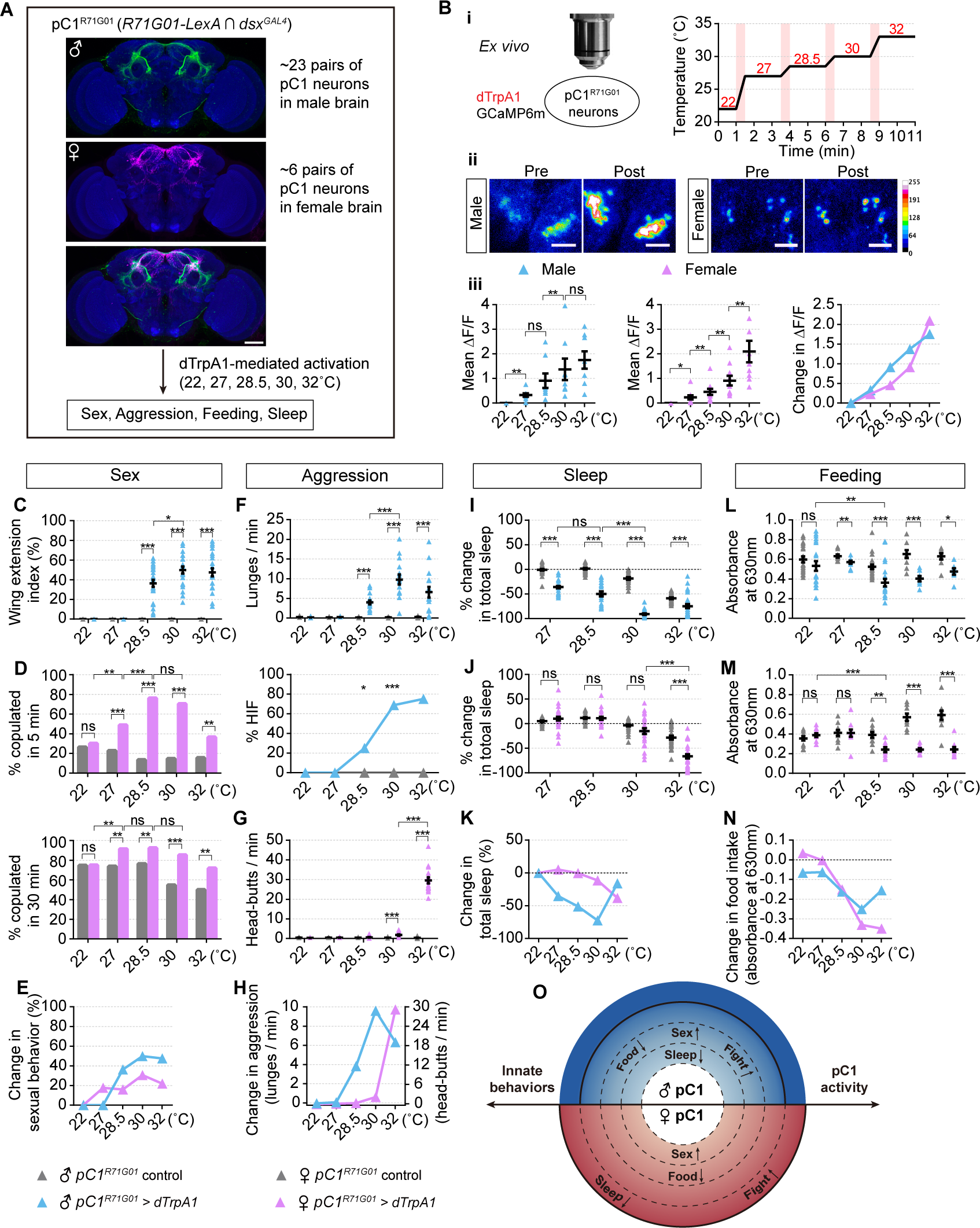
Sexually dimorphic pC1 neurons hierarchically control multiple innate behaviors in both sexes. (A) Overview of the experimental design for the effects of pC1^R71G01^ neuronal activation mediated by dTrpA1 with intensity titration from 22°C to 32°C on a variety of instinctive behaviors. (B) Temperature-control of dTrpA1-mediated activation levels of pC1^R71G01^ neurons. (i) Illustration of the temperature-control protocol for calcium imaging. GCaMP6m and dTrpA1 were co-expressed in the pC1^R71G01^ neurons. Each pink bar indicates a 30-sec temperature ramping. (ii) Representative pseudo-colored images of pC1^R71G01^ cell bodies with maximum intensity projection of fluorescence before and after stimulation in males and females. Scale bars: 30 μm. (iii) Mean fluorescence changes (ΔF/F) in cell bodies of pC1^R71G01^ neurons for each temperature in males (left) and females (middle). The average values of ΔF/F of males and females at five temperatures were plotted together in the right panel. n=8 and 10 for males and females, respectively. (C-E) pC1^R71G01^ neurons promote sexual behaviors in both sexes. (C) Wing extension index of males in 10 min. (D) Copulation percentage of virgin females within 5 min (top) and 30 min (bottom). (E) Change in sexual behaviors in males and females by subtracting the average behavioral index of the experimental group from the control group at each temperature. For (C), n=21-24 males, and for (D), n=80-96 females. (F-H) pC1^R71G01^ neurons promote aggression in both sexes. (F) Frequency of lunges in male pairs (top) and the fraction of pairs performing high-intensity fights including holding, boxing, and tussling (bottom). (G) Frequency of head-butts in female pairs. (H) Change in aggression in males and females by subtracting the average behavioral index from controls at each temperature. For (F), n=16 male pairs for each, and for (G), n=15-16 female pairs. (I-K) pC1^R71G01^ neurons suppress sleep in both sexes. Percentage of total sleep change compared to 22°C baseline in males (I) and females (J). (K) Change in sleep in males and females by subtracting the average behavioral index from controls at each temperature. For (I), n=21-32 males and for (J), n=25-31 females. (L-N) pC1^R71G01^ neurons suppress feeding in both sexes. The amount of ingested food characterized by absorbance of blue dye extraction from the whole flies at 630nm in males (L) and females (M). (N) Change in food intake in males and females by subtracting the average behavioral index from controls at each temperature. n=7-22 tests in males and n=9-12 tests in females, 10 flies were used for each test. (O) A proposed sexually dimorphic hierarchical model for multiple innate behaviors. See Table S1 for detailed statistical comparisons in this and all subsequent figures. ns, not significant, ^∗^p<0.05, ^∗∗^p<0.01, ^∗∗∗^p<0.001, error bars indicate SEM. The full genotypes of flies and sample sizes are also listed in Table S1. See also Figure S1.

We first tested how different pC1^R71G01^ activity levels would control ‘qualitatively’ dimorphic behaviors, sex and aggression, in the two sexes. We found that wing extension, indicative of male courtship, was initiated by a solitary male fly with moderate activation at 28.5℃ and further increased at 30℃, meanwhile, reached saturation (Figure 1C). A similar phenotype was observed in male aggression, where inter-male lunges as well as the fraction of pairs performing high intensity fight (HIF), including holding, boxing, and tussling, were promoted at 28.5℃ and reached to the maximum at 30℃ (Figure 1F). By contrast in females, sexual receptivity was promoted at the minimum stimulation intensity (27℃) and further increased but saturated at 28.5℃ (Figure 1D, 1E and S1A), whereas the frequency of female-specific aggression — head-butts was slightly induced at 30℃ and facilitated remarkably at the highest temperature (32℃) (Figure 1G and 1H). These results show that female pC1^R71G01^ neurons hierarchically promote sexual receptivity and female-specific aggression in an activity-dependent manner, while male pC1^R71G01^ neurons elicit courtship and male-specific aggression with similar thresholds.

We next assayed how pC1^R71G01^ activity would control ‘quantitatively’ dimorphic behaviors, sleep and feeding, in males and females. We found that male flies slept less with the minimum activation of pC1^R71G01^ neurons at 27℃, and such sleep loss reached to the maximum at 30℃ (Figure 1I and S1B). In contrast, female sleep was only suppressed with the strongest activation at 32℃ (Figure 1J, 1K and S1C). To test whether activating pC1^R71G01^ neurons caused a sexual difference in feeding control, we starved flies for 24 hours and assayed their feeding by quantifying the ingested blue dye food which contains both sucrose and yeast at different temperatures. We found that both male and female flies reduced feeding with moderate pC1^R71G01^ activation at 28.5℃ (Figure 1L-1N). Hence, our data indicate that pC1^R71G01^ neurons control sleep differently and feeding similarly in the two sexes.

In summary, our titrated stimulation experiments reveal that homologous but sexually dimorphic pC1^R71G01^ neurons modulated multiple innate behaviors in a sex-specific and activity-dependent manner. Based on these data, we propose a hierarchical model involving a sexually dimorphic center (pC1^R71G01^) that prioritizes distinct behaviors differently in the two sexes (Figure 1O). As pC1^R71G01^ activity rises, males would first reduce sleep, thereby increasing wakefulness, and then elicit male courtship and aggression, while also inhibiting feeding with moderate neuronal activity. By contrast in females, increasing pC1^R71G01^ activity would first promote female receptivity, then inhibit feeding, and lastly induce fighting and suppress sleep.

### Arousal states induced by pC1 neurons are sexually dimorphic

We noticed that the most significant difference between males and females by activating pC1^R71G01^ neurons was that sleep suppression required the minimum activation of pC1^R71G01^ neurons in males but the strongest activation in females. These could be due to general locomotor differences in the two sexes. Thus, we additionally used videos to track the spontaneous locomotor activity of individual flies for 24 hours. Indeed, we found that the daily average velocity of spontaneous walking was significantly increased with the minimum activation at 27℃ and reached to its maximum at 30℃ in males (Figure 2A and 2C). By contrast in females, a slight decrease of locomotion was first observed with moderate activation at 28.5℃, and hyperactivity was only induced with the strongest activation at 32℃ (Figure 2B, 2D and 2E). Thus, the spontaneous locomotor control mediated by pC1^R71G01^ neurons, like sleep control, was also sexually dimorphic.

**Figure 2.**
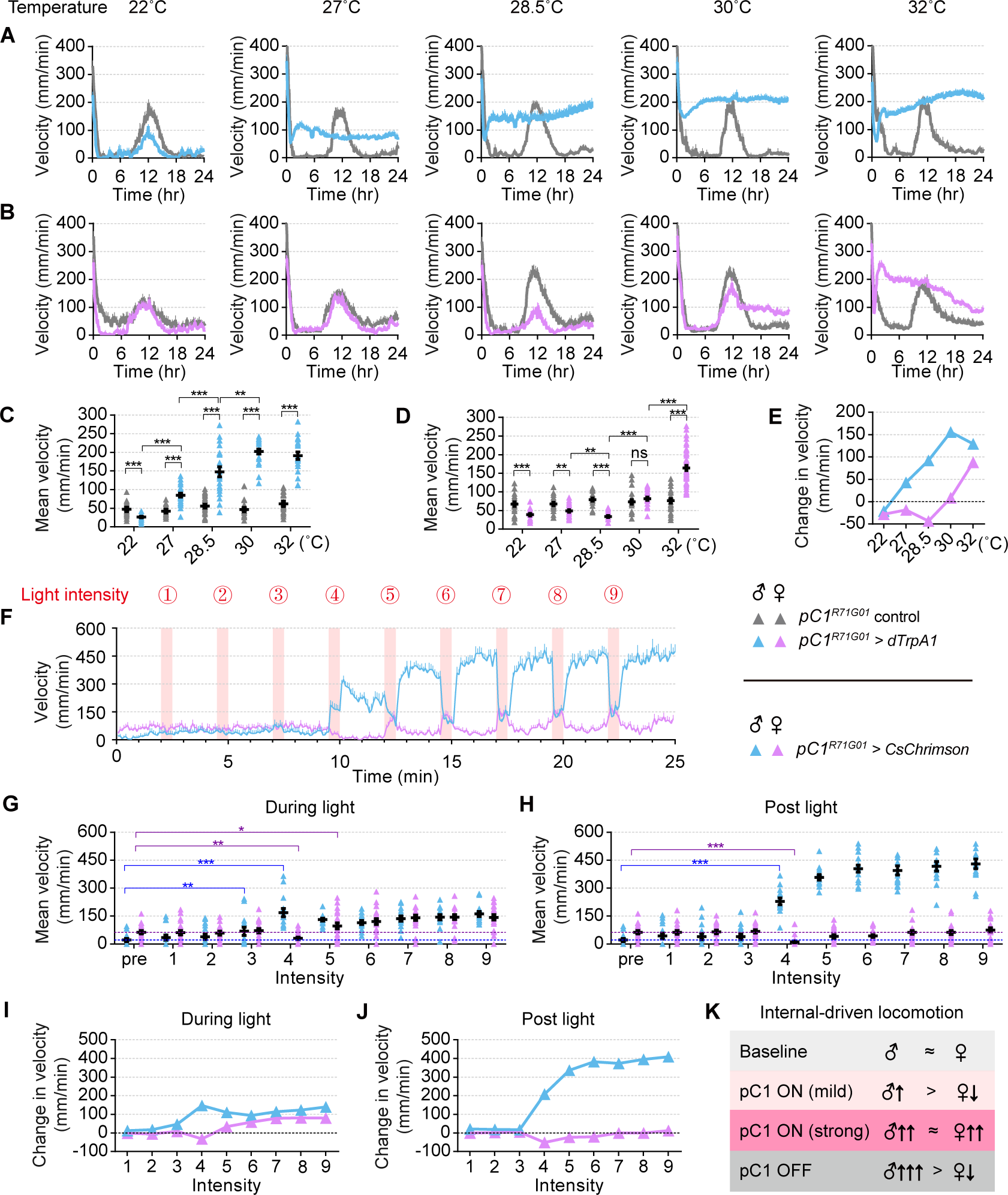
pC1-mediated locomotion patterns are sexually dimorphic. (A-E) Effects of thermogenetic stimulation of pC1^R71G01^ neurons on locomotor activity in both sexes. 24-hour locomotion profiles of males (A) and females (B) during thermogenetic activation of pC1^R71G01^ neurons at different temperatures, which were quantified as daily mean velocity in (C) and (D) respectively. (E) Change in velocity in males and females by subtracting the daily mean velocity of the experimental group from the control group at each temperature. n=20-48 males and n=20-47 females. (F-J) Effects of optogenetic stimulation of pC1^R71G01^ neurons on locomotor activity in both sexes. (F) The average velocity per 5 seconds for both sexes of *pC1^R71G01^>CsChrimson* flies during an intensity titration experiment. Each pink bar indicates a 30-sec photostimulation with increased illumination intensities of a constant red light. (G and H) The quantification of mean velocity per minute for both sexes of *pC1^R71G01^>CsChrimson* flies during (G) and after (H) photostimulation. (I and J) Change in velocity in males and females by subtracting the mean velocity at the baseline phase from the one at each experimental phase. The first two minutes of the experiment represent the baseline phase. n=15 males and n=21 females. (K) Summary of sexually dimorphic locomotion patterns induced by pC1^R71G01^ neurons.

Such sexually dimorphic locomotor control was further proved by optogenetic experiments. Due to the lower precisions in the time resolution and intensity domain of thermogenetic protocols, we drove the expression of red-shifted channel CsChrimson in pC1^R71G01^ neurons for photostimulation. We activated individual *pC1^R71G01^>CsChrimson* flies from both sexes using a constant red light with nine increasing intensities (level 1 to 9, Figure 2F). Consistently, during the light-on phase, there was a gradual enhancement in male locomotor activity as the stimulation intensity rose from level 3 to 4, but higher photostimulation intensities in the later period no longer increased it. On the contrary, at the intensity that significantly promoted male locomotion, females’ movement was inhibited instead, indicating opposite effects by moderate activation of pC1^R71G01^ neurons, which is also consistent with dTrpA1-mediated activation. Surprisingly, as the intensities continued to increase, the locomotor activity of female flies was promoted to a level comparable to that of males (Figure 2G and 2I). Furthermore, another obvious difference between the sexes appeared in the 2-min phase after photostimulation. We observed a more remarkably enhancement of male locomotion following the light offset, indicative of a persistently increased internal state. Nevertheless, female locomotion returned to baseline levels soon after photostimulation (Figure 2H, 2J and 2K). Note that the data we tracked here for only 2 min after light did not exclude the possible existence of a persistently increased internal state upon pC1^R71G01^ activation in females. Taken together, these thermogenetic and optogenetic experiments clearly demonstrate that arousal states induced by pC1 neurons are sexually dimorphic.

### Identification of two distinct types of pC1 neurons controlling locomotor activity oppositely in the two sexes

It has been found that female pC1 neurons could be divided into subtypes at the single-cell level[19], and different pC1 subtypes regulated sexual and aggressive behaviors, respectively[17, 19, 20]. We propose that distinct pC1 subsets may have different activation thresholds to hierarchically control multiple behaviors. Indeed, our calcium tracing results confirmed that female pC1 neurons were functionally heterogeneous and displayed different excitability (Figure S2). However, much less is known for pC1 subsets, especially molecularly defined subtypes, in males.

To identify molecular markers for pC1 neurons, we isolated male pC1 neurons for RNA-sequencing and found a neuropeptide-encoding gene *DH44* was highly expressed in these cells (Figure 3A). Neuropeptide DH44 is the fly homolog of mammalian corticotropin-releasing factor (CRF) and was identified to be expressed dominantly in the pars intercerebralis (PI) region of the brain[35–38]. To verify if *DH44* was expressed in *dsx* neurons, we first performed intersectional analysis of the two knock-in drivers, *dsx^GAL4^* and *DH44^LexA^*, and observed a cluster of neurons located in the pC1 area in both sexes, which labeled ∼6 pairs in male brain and ∼2 pairs in female brain (Figure 3B and 3G). Double staining with anti-DH44 antibody confirmed that at least four out of the six labeled male cells and the two female cells indeed co-localized with DH44 (Figure 3C and 3H). We therefore referred to these neurons existing in both sexes as pC1^DH44^. Similar pC1 neurons also appeared to be tagged by a reverse intersection using *DH44^GAL4^*and *dsx^LexA^*, which additionally targeted pC2 neurons, nevertheless (Figure S3A and S3B). We additionally generated a knock-in *DH44^AD^*line and combined with *dsx^DBD^* as split-GAL4, which also labeled pC1 neurons in both sexes (referred to as pC1^DH44-SS^, Figure S3C and S3D). Together, these findings identify a subset of *dsx*-positive pC1 neurons that also express *DH44*, and these pC1^DH44^ neurons are homologous but sexually dimorphic.

**Figure 3.**
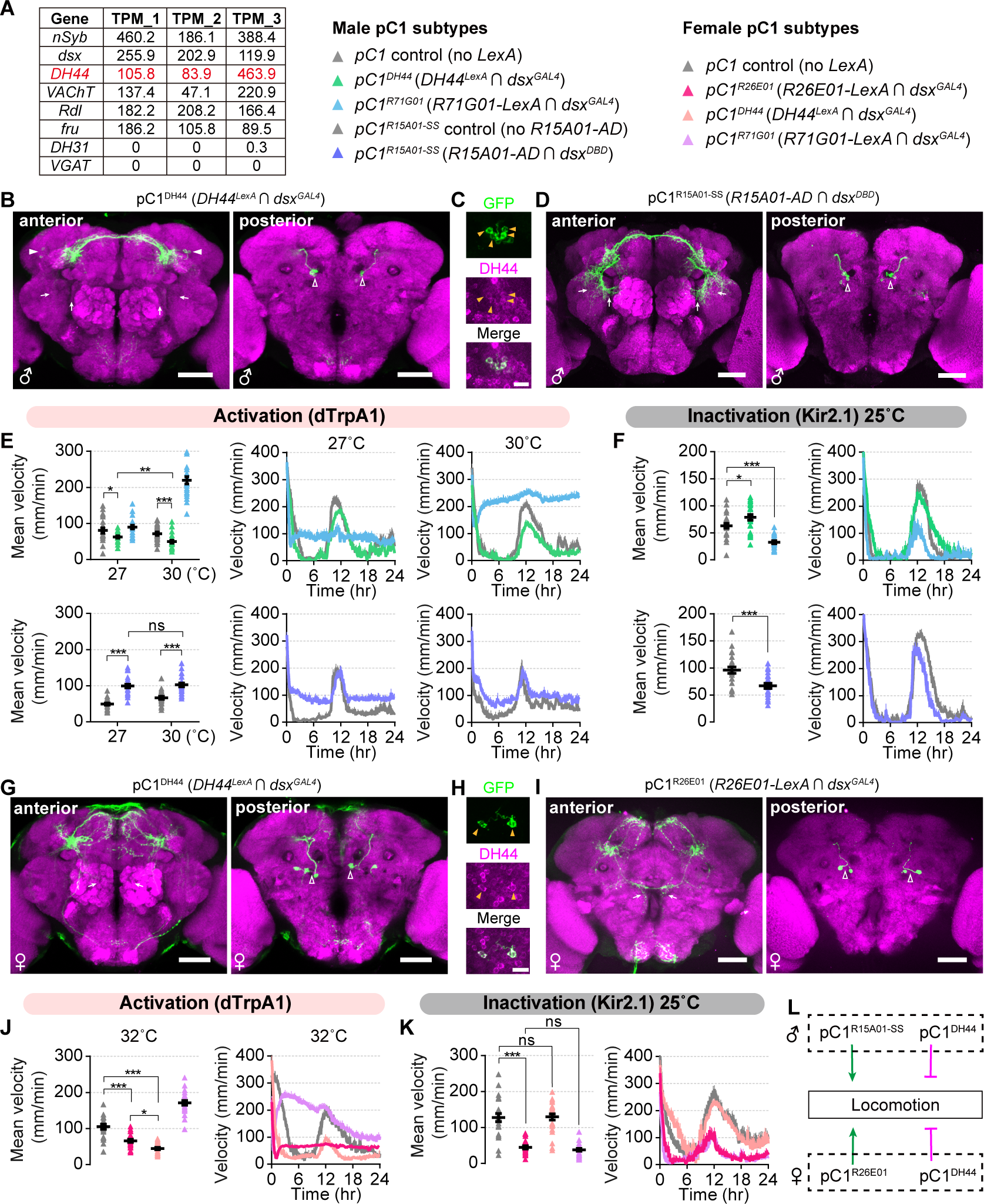
Identification of two distinct types of pC1 neurons controlling locomotor activity oppositely in the two sexes. (A) Representative gene expression in male pC1 neurons from RNA sequencing. Three replicates were performed. (B) Confocal images showing anterior (left) and posterior (right) parts of pC1^DH44^ neurons in the *UAS-FRT-stop-FRT-CsChrimson/+; LexAop-FLP, dsx^GAL4^*/*DH44^LexA^* male brain, co-stained with anti-GFP (green) and anti-nc82 (magenta). Scale bars: 50 μm. (C) Co-localization of pC1^DH44^ neurons and neuropeptide DH44 in the *UAS-FRT-stop-FRT-CsChrimson/+; LexAop-FLP, dsx^GAL4^*/*DH44^LexA^* male brain, co-stained with anti-GFP (green) and anti-DH44 (magenta). Solid yellow arrowheads denote DH44-positive cells. Scale bars: 10 μm. (D) Confocal images showing anterior (left) and posterior (right) parts of pC1^R15A01-SS^ neurons in the *R15A01-AD*/*UAS-myrGFP*; *dsx^DBD^*/*+* male brain, co-stained with anti-GFP (green) and anti-nc82 (magenta). Scale bars: 50 μm. (E and F) 24-hour locomotion profile and mean velocity for pC1^DH44^, pC1^R71G01^ and pC1^R15A01-SS^ activation (E) and silencing (F) experiments in males. n=22-24. (G) Confocal images showing anterior (left) and posterior (right) parts of pC1^DH44^ neurons in the *UAS-FRT-stop-FRT-CsChrimson/+; LexAop-FLP, dsx^GAL4^*/*DH44^LexA^* female brain, co-stained with anti-GFP (green) and anti-nc82 (magenta). Scale bars: 50 μm. (H) Co-localization of pC1^DH44^ neurons and neuropeptide DH44 in the *UAS-FRT-stop-FRT-CsChrimson/+; LexAop-FLP, dsx^GAL4^*/*DH44^LexA^* female brain, co-stained with anti-GFP (green) and anti-DH44 (magenta). Solid yellow arrowheads denote DH44-positive cells. Scale bars: 10 μm. (I) Confocal images showing anterior (left) and posterior (right) parts of pC1^R26E01^ neurons in the *UAS-FRT-stop-FRT-CsChrimson/R26E01-LexA; LexAop-FLP, dsx^GAL4^*/*+* female brain, co-stained with anti-GFP (green) and anti-nc82 (magenta). Scale bars: 50 μm. (J and K) 24-hour locomotion profile and mean velocity for pC1^DH44^, pC1^R26E01^ and pC1^R71G01^ activation (J) and silencing (K) experiments in females. n=22-24. (L) A summary of distinct pC1 subsets oppositely regulating locomotion in both sexes. For (B, D, G and I), open arrowheads denote cell bodies of pC1 subsets; filled arrowheads denote cell bodies of aDN neurons; small arrows compare the projection differences between pC1 subsets. Temperatures for behavioral tests were indicated above each panel. See also Figure S2-4.

Previously, ∼10 pairs of male-specific P1^a^ neurons labeled by *R15A01-AD* and *R71G01-DBD*, some of which were *dsx*-positive (see below), as well as ∼2 pairs of female-specific pC1 neurons by *R26E01-AD* and *dsx^DBD^* (referred to as pC1^R26E01-SS^, Figure S3F and S3H) were found to be responsible for hyperaggression in both sexes[11, 15]. Additionally, we performed intersectional analysis of *dsx^GAL4^*with *R15A01-LexA* or *R26E01-LexA* for comparable labeling with the above pC1^R71G01^ and pC1^DH44^ neurons. Intersection of *dsx^GAL4^*and *R15A01-LexA* labeled both pC1 and non-pC1 neurons in males (Figure S3E), while intersection of *dsx^GAL4^* and *R26E01-LexA* successfully labeled ∼2 pairs of pC1^R26E01^ neurons in females (Figure 3I). We consequently explored the expression patterns using *R15A01-AD* and *dsx^DBD^*, and observed 5-7 pairs of male-specific pC1 neurons (referred to as pC1^R15A01-SS^, Figure 3D and S3G). Male pC1^DH44^ and pC1^R15A01-SS^ neurons displayed morphological differences, in which the ring-shaped arborization was only obvious in pC1^R15A01-SS^ but not pC1^DH44^ neurons. Further intersection between *pC1^R15A01-SS^* and *DH44^LexA^* labeled one neuron per brain, indicating that these were largely distinct pC1 subpopulations. Moreover, female pC1^R26E01^ but not pC1^DH44^ neurons sent long projections to the ventromedial region. Intersection of *pC1^R26E01-SS^* with *DH44^LexA^*also labelled one cell per brain, whose morphology resembled that of the pC1c neuron. Additive expression of *pC1^R26E01-SS^* and *pC1^DH44-SS^* further confirmed their distinct expression yet still with a common cell (Figure S3I-S3N). Thus, we subdivided pC1 neurons in both sexes into a class of sexually dimorphic pC1^DH44^ neurons and a class of sex-specific neurons (male pC1^R15A01-SS^ and female pC1^R26E01-SS^).

We next sought to investigate the effects of these pC1 subtypes on innate behaviors and began by examining the locomotor activity. Remarkably, dTrpA1-mediated activation of pC1^DH44^ neurons at 27℃ decreased male locomotion whereas activation of pC1^R15A01-SS^ neurons increased it. Furthermore, stronger activation of pC1^R71G01^ and pC1^DH44^, but not pC1^R15A01-SS^ neurons, at 30℃ induced more severe effects in locomotor activity (Figure 3E and S4A). Consistent with the above activation experiments, inactivation of pC1 neurons by expressing the inwardly rectifying potassium channel Kir2.1, which caused hyperpolarization of neurons, led to opposite results (Figure 3F). Together, our data reveal that pC1^DH44^ neurons suppress locomotion while pC1^R15A01-SS^ neurons promote locomotion in males.

In females, we used the same intersectional strategy to manipulate pC1^R71G01^, pC1^DH44^, and pC1^R26E01^ neurons to make the results more comparable. Like in males, thermogenetic activation of pC1^DH44^ neurons decreased female locomotion over a 24-h period. In contrast, activating pC1^R26E01^ neurons altered the locomotor pattern by decreasing locomotion in the morning and evening peaks but increasing locomotion elsewhere (Figure 3J and S4D). Moreover, inactivation of pC1^R26E01^ and pC1^R71G01^ decreased female locomotion, but inactivation of pC1^DH44^ neurons had no impact on it (Figure 3K). These results suggest that pC1^DH44^ neurons suppress locomotion and pC1^R26E01^ neurons generally promote locomotion in females. We propose that the lower speed mediated by pC1^R26E01^ neurons might be due to the presence of a DH44-positive cell in these neurons. It is worth noting that the extreme hyperactivity induced by pC1^R71G01^ neurons in both sexes was not repeated in any of the subtypes.

In summary, we certify that pC1 neurons comprise different populations with heterogeneous molecular expression and identify two types of pC1 neurons in each sex to oppositely control locomotor activity (Figure 3I).

### pC1 subpopulations in both sexes regulate specific innate behaviors

We then examined whether these two pC1 subpopulations described above in each sex regulated other innate behaviors. Strikingly, dTrpA1-mediated activation of pC1^DH44^ and pC1^R15A01-SS^ subtypes failed to enhance male wing extension or inter-male aggression as pC1^R71G01^ activation did (Figure 4A, 4B and S4B). Meanwhile, silencing pC1^R15A01-SS^ neurons did not impair either courtship intensity or mating success, however, silencing pC1^DH44^ neurons decreased the courtship index and the rate of successful copulation, although not as severe as silencing pC1^R71G01^ neurons (Figure 4D). Besides, silencing either pC1^DH44^ or pC1^R15A01-SS^ neurons did not affect inter-male aggression (Figure 4E). As for feeding, activating or silencing pC1^DH44^, like manipulating pC1^R71G01^ neurons, both resulted in a lower amount of ingested food. On the other hand, activation of pC1^R15A01-SS^ neurons suppressed feeding and silencing them promoted food intake (Figure 4C, 4F and S4C). These results reveal that pC1^R71G01^ and pC1^DH44^ neurons have U-shaped control in feeding, while pC1^R15A01-SS^ neurons generally inhibit feeding in males.

**Figure 4.**
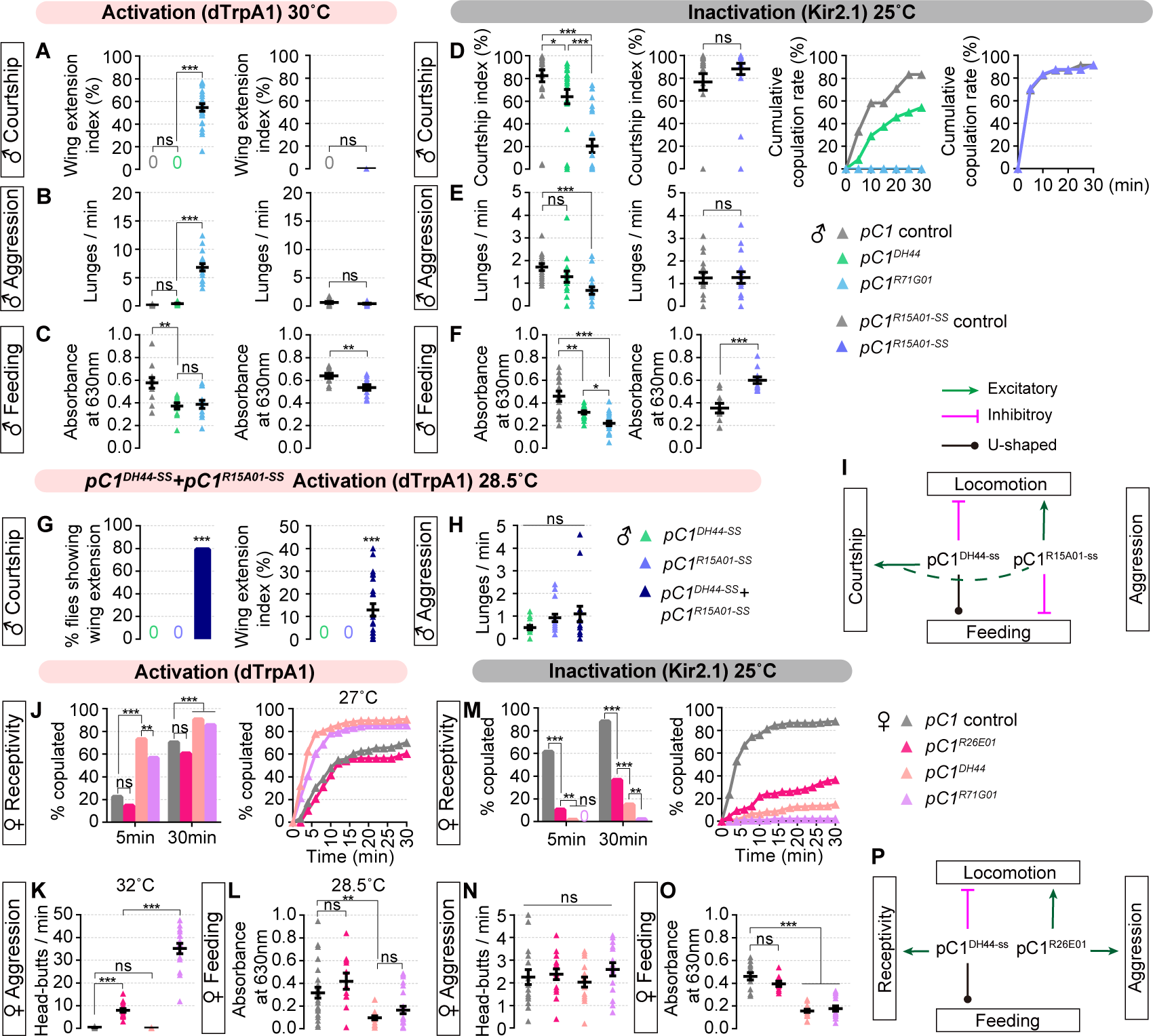
pC1 subpopulations regulate distinct behaviors parallelly and/or collectively in a sexually dimorphic way. (A-C) Wing extension index (A), frequency of lunges (B), and food intake characterized by absorbance at 630nm (C) for pC1^DH44^, pC1^R71G01^ and pC1^R15A01-SS^ activation experiments in males. For (A), n=24 males for each. For (B), n=12-16 male pairs. For (C), n=10-12 tests, and each test used 10 flies. (D-F) Courtship index in 10 min and cumulative copulation rate within 30 min (D), frequency of lunges (E), and food intake characterized by absorbance at 630nm (F) for pC1^DH44^, pC1^R71G01^ and pC1^R15A01-SS^ inactivation experiments in males. For (D), n=23-24 males. For (E), n=14-16 male pairs. For (F), n=10-20 tests, and 10 flies were used for each test. (G and H) Wing extension performance (G) and frequency of lunges (H) for joint activation of pC1^DH44-SS^ and pC1^R15A01-SS^ neurons in males. For (G), n=24 males for each. For (H), n=15-16 male pairs. (I) Summary of male pC1 subsets regulating sex, aggression, feeding and locomotion. (J-L) Copulation percentage (J), frequency of head-butts (K), and food intake characterized by absorbance at 630nm (L) for pC1^DH44^, pC1^R26E01^ and pC1^R71G01^ activation experiments in females. For (J), n=48-144 females. For (K), n=15-16 female pairs. For (L), n=11-25 tests, and 10 flies were used for each test. (M-O) Copulation percentage (M), frequency of head-butts (M), and food intake characterized by absorbance at 630nm (O) for pC1^DH44^, pC1^R26E01^ and pC1^R71G01^ inactivation experiments in females. For (M), n=96-160 females. For (N), n=16 female pairs for each. For (O), n=9-17 tests, and 10 flies were used for each test. (P) Summary of female pC1 subsets regulating sex, aggression, feeding and locomotion. Magenta lines indicate inhibitory functions. Green lines indicate excitatory functions. Dashed green curve represents a synergistic function from pC1^R15A01-ss^ to pC1^DH44^ neurons. Black lines with dots represent U-shaped functions, in which both activation and inactivation lead to a reduction of feeding. Temperatures for behavioral tests were indicated above each panel. See also Figure S4.

The inability of pC1 subpopulations to induce courtship or aggression compared with pC1^R71G01^ in males raised the possibility that there were additional subtypes responsible for these behaviors, or there was synergetic function from pC1^DH44^ and pC1^R15A01-SS^ neurons. To determine that, we sought to unite pC1^DH44-SS^ and pC1^R15A01-SS^ and jointly activate them. Combining the *DH44^AD^* and *R15A01-AD* with *dsx^DBD^*resulted in a markedly increased number of cells in the pC1 area (Figure S3O). We found that such combinational activation of the two pC1 subpopulations was sufficient to induce wing extension (Figure 4G), which is consistent with previous findings that a larger number of pC1 neurons were required to induce wing extension [11, 22]. Co-activation of the two pC1 subpopulations failed to stimulate inter-male aggression (Figure 4H). Together these results demonstrate distinct roles of pC1^DH44-SS^ and pC1^R15A01-SS^ neurons in regulating male locomotor, courtship and feeding behaviors (Figure 4I).

In females, both activation and inactivation experiments showed that pC1^DH44^ subpopulations, similar to pC1^R71G01^ neurons, were crucial for virgin female receptivity (Figure 4J, 4M and S4E). Although pC1^R26E01^ neurons were insufficient to promote receptivity, silencing such neurons resulted in decreased receptivity, possibly because one of the cells was DH44-positive. In contrast, activating pC1^R26E01^ but not pC1^DH44^ subpopulations induced inter-female aggression, although silencing these neurons did not affect the already low level of female aggression (Figure 4K, 4N and S4F). Additionally, pC1^DH44^, like pC1^R71G01^ neurons, were involved in the regulation of feeding, with their activation or suppression reducing the food intake, but either activating or silencing pC1^R26E01^ neurons did not affect it (Figure 4L, 4O and S4G). These results indicate that female pC1^DH44^ and pC1^R26E01^ neurons have differential roles in mediating female receptivity, aggression, feeding and locomotor behaviors (Figure 4P).

In summary, pC1 subpopulations in both sexes control specific innate behaviors. Synergetic effects are observed in male but not female pC1 subsets to regulate sexual behaviors. These results also suggest the existence of an unidentified male pC1 subset for promoting aggression.

### Neuropeptide DH44 is crucial for sexually dimorphic locomotor activity control

The aforesaid results raised several questions, such as how pC1^DH44^ neurons regulate multiple behaviors and what role the neuropeptide DH44 acts in the modulation of these behaviors. In addition, the major fast excitatory neurotransmitter acetylcholine (ACh) was proved to be produced by the majority of pC1 neurons in both sexes[10]. Therefore, we sought to identify the signals required by pC1 neurons using RNAi against *DH44* and/or *choline acetyltransferase* (*ChAT*), which encodes a biosynthetic enzyme for ACh. The effectiveness of the *UAS-DH44IR* line was verified by the lethal effect during the pupal stage under the control of *actin-GAL4* and the negative immunostaining signal with anti-DH44 using the *DH44^GAL4^* driver (Figure S5A and S5B). The effectiveness of the *UAS-ChATIR* line was verified by the lethal effect under the control of *actin-GAL4* and used previously[39].

We first investigated what role did ACh play in locomotion modulation. RNAi-mediated knockdown of *ChAT* consistently reduced locomotor activity in both sexes, regardless of driven by *dsx^GAL4^* (Figure 5A and 5B) or pC1 subsets (Figure S5C, S5E, S5G and S5H) under physiological states, or with specific pC1 neurons being activated via dTrpA1 (Figure 5C, 5D, 5K, S5D, S5F, S5I and S5J). We next investigated the role of neuropeptide DH44. Surprisingly, knocking down *DH44* in *dsx* neurons increased male locomotion but rarely influenced female locomotion (Figure 5A and 5B), which is consistent to the results of silencing pC1^DH44^ neurons (Figure 3F and 3K). A similar result was obtained from restricting the RNAi-mediated knockdown of *DH44* to pC1^DH44-SS^ neurons (Figure S5G and S5H). Since activation of pC1^DH44^ neurons reduced locomotor activity in both sexes, we continued to explore whether this process would act through DH44. We knocked down *DH44* in pC1^DH44-SS^ neurons while activating them and indeed observed a slight raise in males and an obvious recovery in females relative to the activation of pC1^DH44-SS^ neurons alone, nevertheless, the male daily mean velocity was not significant (Figure 5C and 5D). However, analysis of the mean velocity during a period from ZT1 to ZT5 supported a significant recovery of locomotion in males with *DH44* knockdown (Figure 5C). Thus, quantifying increases in the velocity of *DH44*-knockdown flies from both sexes reveals that pC1^DH44^ neurons exert the function of inhibiting locomotion through neuropeptide DH44. Moreover, double knockdown of *DH44* and *ChAT* suggested an additive effect of DH44 and ACh in locomotor control under most conditions (Figure 5D, S5G, S5H and S5J); while in cases of male pC1 activation, the loss of both neuromodulators reduced velocity to the similar levels as the loss of ACh alone, suggesting a converged output by ACh (Figure 5C and S5I).

**Figure 5.**
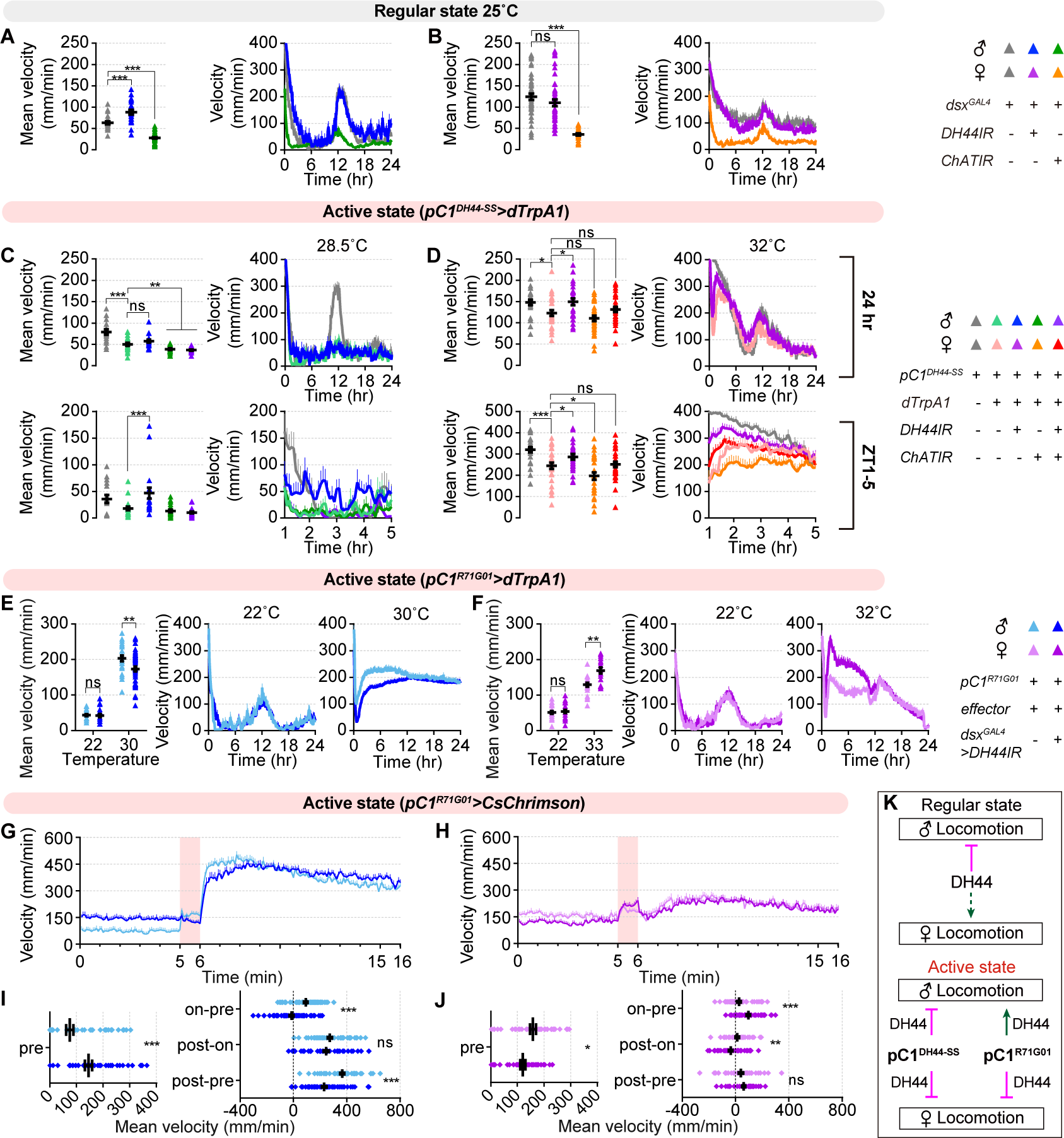
Neuropeptide DH44 is crucial for sexually dimorphic locomotor activity control. (A and B) 24-hour locomotion profile and mean velocity with *DH44* or *ChAT* knockdown in *dsx^GAL4^* neurons in males (A) and females (B). n=23-45 for males and n=23-46 for females. (C and D) 24-hour (top) and 4-hour (bottom) locomotion profile and mean velocity for males (C) and females (D) during activation of pC1^DH44-SS^ neurons with *ChAT*, *DH44*, or both knockdown. n=16-20 for males and n=24-30 for females. (E and F) 24-hour locomotion profile and mean velocity for males (E) and females (F) during dTrpA1-mediated thermoactivation of pC1^R71G01^ neurons with *DH44* knockdown in *dsx^GAL4^* neurons. n=24-41 for males and n=18-24 for females. (G-J) Locomotor activity for males (G) and females (H) during CsChrimson-mediated photoactivation of pC1^R71G01^ neurons with *DH44* knockdown in *dsx^GAL4^* neurons, which were quantified in (I) and (J), respectively. Each pink bar indicates a 1-min photostimulation. (I, J) Mean velocity in 5 minutes before photostimulation (left), and velocity changes during or after photostimulation (right). 2 minutes post photostimulation was used for analysis. For (G and I), n=58 males for each and for (H and J), n=45-47 females. (K) Summary of sex-specific and state-dependent locomotion control by DH44. Magenta lines indicate inhibitory functions. Green lines indicate excitatory functions. Dashed green line represents a weak stimulatory function from (J, baseline). Temperatures for behavioral tests were indicated above each panel. See also Figure S5.

Given that activating pC1^R71G01^ neurons dramatically increased locomotion in both sexes, we wondered how DH44 was involved. To our surprise, after knocking down *DH44* in *dsx*-positive neurons at a state of activating pC1^R71G01^ neurons with high intensity, male flies locomoted much more slowly while female flies locomoted much faster (Figure 5E and 5F). These data reveal that the already high locomotor activity resulting from intense activation of pC1^R71G01^ neurons is still suppressed by DH44 in females but promoted by DH44 in males. Hence, DH44 may oppositely regulate locomotor activity in males using a state-dependent manner. To further confirm this, we took advantage of the optogenetics, where the higher precisions in the time resolution can allow us to detect this state-dependent effect more rapidly. After a 5-min baseline recording, we optogenetically activated pC1^R71G01^ neurons for 1 min in a solitary male or female fly, and continued to record locomotion for 10 min after light went off (Figure 5G and 5H). As expected, *pC1^R71G01^>CsChrimson* males showed a robust increase in locomotor activity during photostimulation and went much higher after light offset. Comparatively, males with additional *dsx*-driven knockdown of *DH44* showed higher activity in baseline but no enhancement during photostimulation stage and a much less enhancement during the 2-min stage after photostimulation (Figure 5I). In contrast, *pC1^R71G01^>CsChrimson* females moved slightly faster during photostimulation and returned to the baseline levels within a short time after light offset. However, *DH44*-knockdown females with a lower activity in baseline showed a higher locomotor activity during photostimulation and decayed rapidly after photostimulation (Figure 5J). Together, DH44 in males inhibits locomotion in a physiological state and promotes locomotion in an activated state; inversely, DH44 in females inhibits locomotion in an activated state.

In summary, ACh plays an excitatory role to promote locomotion in both sexes while DH44 plays a modulatory role with more complexity, showing both state-dependent and sex-dependent effects (Figure 5K).

### ACh and DH44 orchestrate multiple innate behaviors in both sexes

We subsequently investigated how DH44 and ACh would modulate sexual, aggressive and feeding behaviors. The courtship assay revealed that knockdown of *DH44* in *dsx*-positive neurons attenuated the strength of male courtship as well as the mating success (Figure 6A). These findings are consistent with above results that silencing pC1^DH44^ neurons decreased male courtship (Figure 4D). As shown above, the addition of pC1^DH44-SS^ neurons led to the activation of pC1^R15A01-SS^ neurons to produce a wing extension phenotype in solitary males (Figure 4G), we therefore asked how DH44 and ACh would be involved in this process. We knocked down *DH44* in the unity of pC1^DH44-SS^ and pC1^R15A01-SS^ neurons while activating them, and found that both the fraction of flies performing wing extension and the time that the behavior was displayed were lower than those of the control group, indicating an enhanced role of DH44 (Figure 6B). Additionally, males with the loss of ACh alone or both also showed little wing extension when pC1^DH44-SS^ and pC1^R15A01-SS^ neurons activated combinedly. These results indicate that both DH44 and ACh in pC1 neurons promote male sexual behaviors.

**Figure 6.**
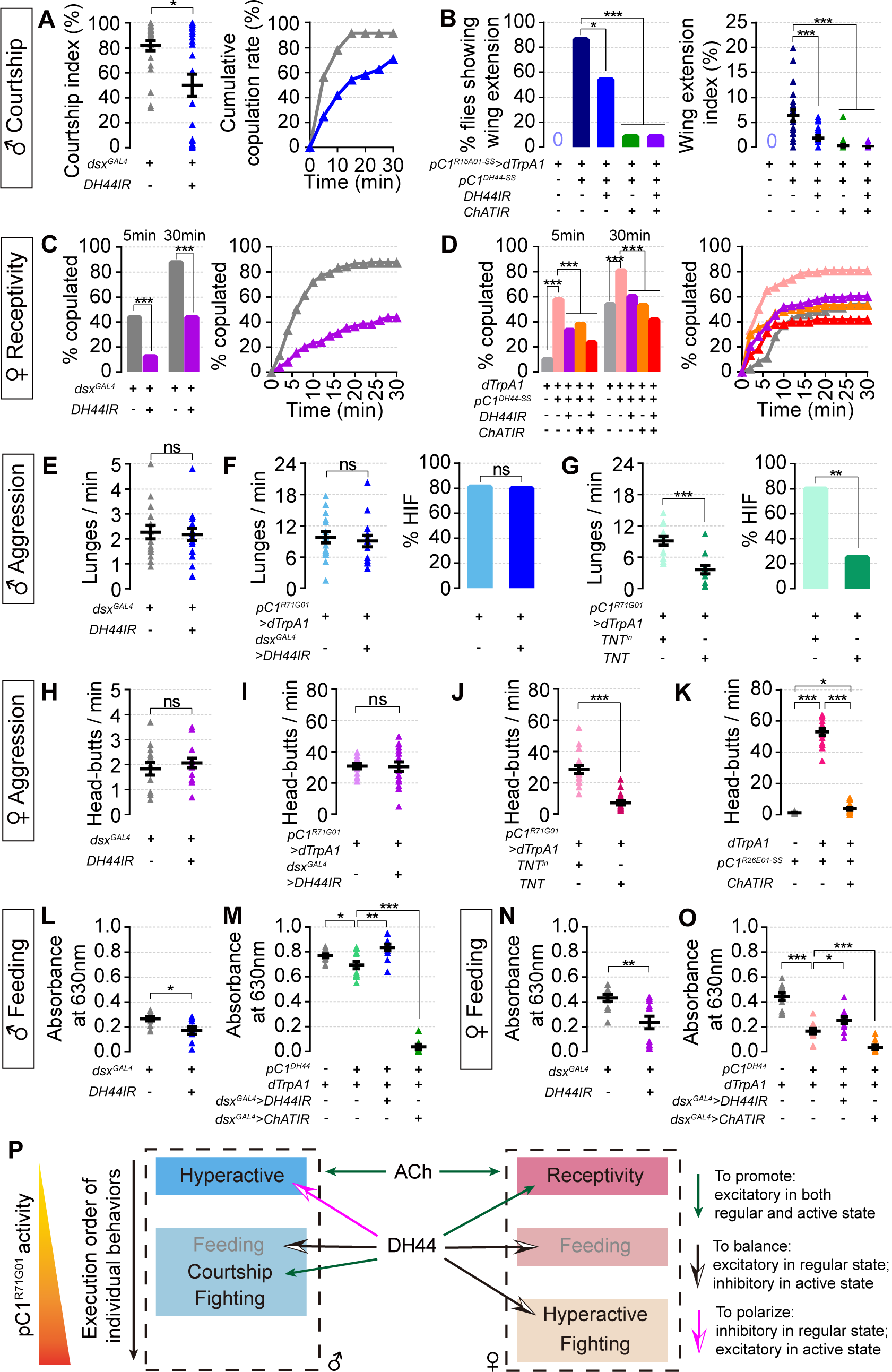
ACh and DH44 orchestrate multiple innate behaviors with different effects in both sexes. (A) Courtship index in 10 min (left) and cumulative copulation rate within 30 min (right) for males with *DH44* knockdown in *dsx^GAL4^* neurons at 25°C. n=24 for each. (B) The fraction of males performing wing extension (left) and wing extension index in 10 min (right) during joint activation of pC1^DH44-SS^ and pC1^R15A01-SS^ neurons with *ChAT*, *DH44*, or both knockdown at 28.5°C. n=22-24. (C) Receptivity curve and copulation percentage within 5 min and 30 min for females with *DH44* knockdown in *dsx^GAL4^*neurons at 25°C. n=96 for each. (D) Receptivity curve and copulation percentage within 5 min and 30 min during activation of female pC1^DH44-SS^ neurons with *ChAT*, *DH44*, or both knockdown at 27°C. n=60-108. (E) Frequency of male lunges with *DH44* knockdown in *dsx^GAL4^* neurons at 25°C. n=16 pairs for each. (F) Frequency of male lunges (left) and the fraction of pairs showing high-intensity fights (right) during activation of pC1^R71G01^ neurons with *DH44* knockdown in *dsx^GAL4^*neurons at 30°C. n=15-16 pairs. (G) Frequency of male lunges (left) and the fraction of pairs showing high-intensity fights (right) during activation of pC1^R71G01^ neurons expressing *TNT* or the inactive control *TNT^in^* at 30°C. n=12-15 pairs. (H) Frequency of female head-butts with *DH44* knockdown in *dsx^GAL4^* neurons at 25°C. n=14-15 pairs. (I) Frequency of female head-butts during activation of pC1^R71G01^ neurons with *DH44* knockdown in *dsx^GAL4^* neurons at 32°C. n=15-16 pairs. (J) Frequency of female head-butts during activation of pC1^R71G01^ neurons expressing *TNT* or *TNT^in^* at 32°C. n=16 pairs for each. (K) Frequency of female head-butts during activation of pC1^R26E01-SS^ neurons with *ChAT* knockdown at 32°C. n=16 pairs for each. (L and N) Food intake characterized by absorbance at 630nm for males (L) and females (N) with *DH44* knockdown in *dsx^GAL4^* neurons at 25°C. n=11 tests in males for each and n=11-12 tests in females, 10 flies were used for each test. (M and O) Food intake characterized by absorbance at 630nm for males (M) and females (O) during activation of pC1^DH44^ neurons with *DH44* or *ChAT* knockdown in *dsx^GAL4^* neurons at 28.5°C. n=8-11 tests in males and n=10-12 tests in females, 10 flies were used for each test. (P) Summary of different roles of ACh and DH44 in regulating sex, aggression, feeding and locomotion in males and females. Grey for ‘feeding’ indicates suppression of the behavior, in contrast to promotion of other behaviors, via pC1 activation. See also Figure S6.

On the other hand, in the female reception assay, we found that knocking down *DH44* driven by *dsx^GAL4^* (Figure 6C), or *pC1-SS2* (Figure S6A), which targeted five pairs of female pC1 neurons[18], but not *vpoDN-SS1* (Figure S6B), which labeled *dsx* neurons controlling vaginal plate opening (vpo)[19], reduced female receptivity. Furthermore, knockdown of *DH44* or *ChAT* in pC1^DH44-SS^ neurons, regardless of artificial activation of them (Figure 6D) or not (Figure S6C), decreased female receptivity compared to corresponding controls. Interestingly, females that lost ACh in pC1^DH44-SS^ neurons had lower acceptance within 5 min, while females that lost DH44 only began to decrease their receptivity at a later stage (Figure S6C). Such difference may reflect the releasing dynamics of neurotransmitters and neuropeptides, in which transmitters may act more rapidly than peptides. In another genetic setup, knocking down *DH44* in all *dsx* cells while activating pC1^R71G01^ neurons reduced female receptivity to some extent, which was further impaired by expressing tetanus toxin light chain (TNT), a synaptic neurotransmission blocker, but not its inactivation form (TNT^in^) in the pC1^R71G01^ neurons (Figure S6D). Together, these results indicate that both DH44 and ACh in pC1 neurons promote female receptivity and suggest a combinational and dynamic function of them.

As for the capacity of fight in both sexes, however, the frequency of both male lunges and female head-butts did not change after knocking down *DH44* in *dsx* neurons (Figure 6E and 6H). Consistently, the vigorous fighting evoked by activating pC1^R71G01^ neurons was not abrogated by *dsx^GAL4^*–driven knockdown of *DH44* (Figure 6F and 6I). These results reveal that DH44 seems to not participate in the regulation of aggression in either sex. We subsequently suspected neurotransmitters as the primary functional signals. Indeed, expressing TNT in pC1^R71G01^ while activating these neurons reduced aggression in both sexes compared to control flies expressing TNT^in^ (Figure 6G and 6J). Given that pC1^R26E01^ neurons in females were responsible for hyperaggression, therefore, we activated these neurons and knocked down *ChAT* in them. We confirmed that artificial activation of pC1^R26E01-SS^ neurons using split-GAL4 to drive *UAS-dTrpA1* resulted in a strong enhancement of female head-butts. Expectedly, this phenotype was almost completely impaired by knocking down *ChAT* (Figure 6K). Together, these results suggest that ACh but not DH44 in pC1 neurons is crucial for aggressive behaviors in both sexes.

Different in feeding behaviors where both activation and inactivation of pC1 neurons reduced food intake, we thereby exploited the molecular mechanisms briefly. We showed that a loss of DH44 in *dsx* neurons decreased the amount of food intake in both sexes (Figure 6L and 6N), which is consistent with the inhibition of pC1^DH44^ neurons (Figure 4F and 4O). Moreover, knocking down *DH44* in pC1-SS2 neurons resulted in a similar phenotype (Figure S6E). These results suggest that DH44 promotes feeding under physiological conditions in both sexes. But surprisingly, when we conducted intersectional activation of pC1^DH44^ neurons using flies carrying *DH44^LexA^* and *dsx^GAL4^*, a drop in food intake was observed, whereas flies with additional *dsx^GAL4^*-driven *DH44* knockdown at least partially restored the amount of consumed food, indicating that DH44 suppressed feeding under activation conditions. Conversely, flies with *dsx^GAL4^*-driven knockdown of *ChAT* barely eat, supporting the excitatory effect of ACh in feeding (Figure 6M and 6O). These findings demonstrate that ACh in pC1 neurons promotes feeding, while DH44 in pC1 neurons bilaterally regulate it in a state-dependent manner, in both sexes.

In summary, ACh in pC1 neurons promotes locomotor, sexual, aggressive and feeding behaviors in both sexes, while DH44 plays sex-specific and state-dependent roles in mediating sexually dimorphic behaviors (Figure 6P).

## Discussion

Extensive research has provided important insights into the understanding of individual behaviors in a male-biased manner. In this study, we have systematically compared multiple innate behaviors and their prioritizations in the two sexes. We have identified a cluster of sexually dimorphic but homologous pC1 interneurons and a neuropeptide homologous to mammalian CRF functioning for the sexually dimorphic and activity-dependent output of individual behaviors in *Drosophila*. Our finding of the neural basis underlying sexually dimorphic prioritization of behaviors highlights a general strategy to implement both the behavioral adaptations and its sexual dimorphism.

### Sexually dimorphic prioritization of multiple innate behaviors

In most animal species, males devote a lot of time and energy to mating with as many females as possible, therefore, their reproductive success is limited by the ability to access and copulate females, sometimes accompanied by fighting with competitors. Nevertheless, female animals take more time, energy and risk for pregnancy and care, consequently, their reproductive success is limited by the ability to choose better mating partners and spawning grounds. The fact that males and females take on distinct roles in reproduction makes them to select different behavioral strategies, e.g., males behave more actively in a social context (court or fight), while females remain relatively calm for better evaluation and decision-making.

Our study shows a class of homologous pC1 interneurons in the brain of *Drosophila* taking a sexually dimorphic control of multiple behaviors and their prioritization in an activity-dependent manner. With an increasing activity of pC1^R71G01^, males modify their behaviors sequentially by (1) reducing sleep and increasing locomotor activity with minimum activation; and (2) courting or fighting with moderate activation, meanwhile, reducing food intake. In contrast, females respond to the increasing activity of pC1^R71G01^ orderly by (1) being more receptive with mild activation; (2) reducing feeding with stronger activation; and (3) being hyperactive and aggressive with the highest activation. These dimorphic orders of executing behaviors by activating pC1 neurons coincide the sex roles in reproduction by fruit flies: males command active locomotion to attack others for competition or court females with an elaborate ritual, while fussy females are more passive and select to pause when decide to accept[40], yet rarely fight and almost never court. Given that the locomotor activity represents a widely accepted endogenous form of arousal[41, 42], which belongs to internal states with the capacity to change responses to sensory stimulus and lead to behavioral preferences[28, 43], we suggest that the sexual differences in locomotor control may represent dimorphic arousal states in the two sexes, with different scalability and persistence. Indeed, a striking difference in locomotor control from optogenetic activation of pC1^R71G01^ neurons is that males but not females dramatically increase locomotion after photoactivation in a persistent manner. Such dimorphic arousal states controlled by pC1 neurons fit well to the ‘hydraulic’ model by Lorenz of executing different behaviors according to the level of internal states[44].

A similar control in the two sexes by pC1 neurons is to inhibit feeding in a U-shaped manner. We propose that pC1^R71G01^ neurons do not directly control feeding, rather, they appear to affect the normal expression of the feeding module through a crosstalk between survival-related behaviors and reproductive behaviors, resembling a ‘hierarchical’ model of decision-making by Tinbergen[45]. Thus, our results implement both the Lorenz and Tinbergen models and provide a neural mechanism for behavioral priority and its sexual dimorphism, in which dimorphic pC1 neurons serve as a ‘reproductive center’ encoding internal states in both sexes to coordinate different circuits in order and convert the graded multisensory inputs into different motor outputs.

### Redundant and collective control in males vs. centralized control in females

A remarkable feature of sexual dimorphism is that male flies have ∼10 times more pC1 neurons than females, which is not seen or could not be revealed yet due to the technique limit in any other model organism. Despite this difference, both male and female pC1 neurons are heterogeneous. Five morphologically distinct pC1 cells named pC1a-e were found in females benefit from the establishment of an EM database of a female *Drosophila* brain (FAFB)[46], however, the lack of EM volume in a male brain as well as the greater number of male pC1 neurons hindered their classification. Our results here demonstrate that pC1 neurons are molecularly heterogeneous, as only a part of them express the neuropeptide DH44. These DH44-positive subtypes are homologous with different cell numbers in the two sexes, in addition to sex-specific subtypes (pC1^R15A01^ in males and pC1^R26E01^ in females). Silencing any pC1 subset could not eliminate male courtship, indicative of their redundant functions in males. Furthermore, activating the combination of pC1^DH44^ and pC1^R15A01^ neurons, but not either subset alone, induces male wing extension, suggesting their synergistic effects in male courtship. The finding that pC1^R71G01^ neurons, but not the two subsets (pC1^DH44^ or pC1^R15A01^), promote male aggression also suggests the involvement of an unidentified subset of pC1 neurons in aggression, which still awaits further investigation. In contrast, only 1 to 2 out of 6 pairs of female pC1 neurons are crucial for an individual behavior (e.g., sex or aggression)[15–17, 19, 47, 48], which we refer to as a centralized control in contrast to the redundant and collective control in males.

The huge sex differences of the pC1 functions may be causal, but more likely provide some general principles of the organization and operation of sexually dimorphic ‘reproductive centers’ in animals[24]. First, the male reproductive center may contain more neurons with redundant functions to confer courtship robustness to potential but sometimes wrong mates, while fewer counterparts in females with specialized functions make them to be very selective on the quality of stereotypic courtship from males. Second, collective control from distinct subpopulations of neurons in males may generate flexible, scalable, and persistent behavioral outputs, while the centralized control is more decisive such that females either accept or reject males, which also fits the complete transition between female pre-copulatory and post-copulatory behaviors.

### Flexible modulation of co-expressed neuropeptide and neurotransmitter

Our results identify the neuropeptide DH44 as a key regulator, in addition to a classical neurotransmitter ACh, in pC1 and possibly other *dsx* neurons for the sexually dimorphic behavioral prioritization. DH44 has long been thought to be predominantly expressed only in six cells in the pars intercerebralis (PI) region, which is regarded as a neuroendocrine center regulating various physiological processes and behaviors such as fluid secretion[35], stress tolerance[49, 50], rest: activity rhythms[36, 51, 52], brain nutrient sensing[37, 53–55] and female sperm ejection[38]. We find that DH44 also functions in the central pC1 neurons to modulate locomotor, sexual and feeding behaviors in different ways. First, DH44 released from pC1 neurons promotes sexual behaviors in both sexes, the exact mechanism of which still requires further study. Second, DH44 in both sexes promotes feeding physiologically whereas turns to suppress feeding once pC1^DH44^ neurons are activated. Such U-shaped modulation of DH44 has also been found in regulating the rest: activity rhythms[36]. We suspect that this process may be explained by the difference in the amount of DH44 released in physiological and active states as well as the differential sensitivity of its two receptors, DH44-R1 and DH44-R2[56]. Third, while DH44 in pC1 neurons generally inhibits locomotion, it turns to promote the pC1-activation-induced hyperactivity specifically in males. Such a sex-specific and state-dependent modulation of locomotion makes males more active and sensitive to potential mates and/or competitors. Based on the functions of DH44 in both PI and pC1 cells, we propose that DH44 may serve as an internal communicator in the brain evaluating the internal energy status through nutrition sensing and the availability of potential mates to adjust locomotor, feeding and sexual behaviors, which eventually benefit their survival and reproduction.

Synaptic complexity executed by small-molecular transmitters and neuropeptides is well-identified in many neural types through animal kingdom[57, 58]. In contrast to the diverse modulatory functions of DH44, we find that ACh in *dsx* neurons promotes the execution of all tested behaviors in both males and females. ACh may generate fast but brief, ionotropic responses locally and DH44 may induce slow but long-lasting, metabotropic responses in a longer distance. Indeed, we observed complementary roles of ACh and DH44 in female receptivity, where ACh mediates rapid acceptance while DH44 plays a long-term effect. Our results support that the functions of these ‘classical’ and ‘modulatory’ signals can be complementary and non-linear to generate robust yet flexible behavioral outputs in a fixed circuit.

### A general model underlying sexually dimorphic behavioral prioritization

In both invertebrates and vertebrates, a few quantitative differences of the nervous system, especially in the interneurons, are found in the two sexes of the same species establishing the neural basis of sexually dimorphic behaviors[5]. Our findings expand this mechanism by providing a two-layered model that implements both sexual dimorphism and behavioral prioritization across species. The first layer is the pre-determined sexually-dimorphic neural circuits specified by sex determination genes during development[59]. Further, the prioritization of multiple behaviors and its dynamic changes require the second layer at the level of neuromodulation in developed circuits, especially the neuropeptides which are capable to travel a long distance before binding to receptors. In the case of our study, the development of homologous but dimorphic pC1 neurons, mainly specified by *dsx*, establishes the neuronal substrates for sex-specific behaviors. On such a circuit basis, DH44 exerts sex-specific and state-dependent functions to facilitate behavioral prioritization in a sexually dimorphic way, which is also adaptable to internal and external changes.

Our study provides a framework to investigate multiple innate behaviors and their prioritizations in both sexes. Future studies would reveal how DH44 exerts the sex-specific and state-dependent functions in pC1 neurons in molecular details, and whether such a mechanism would apply to high-order animals as counterparts for both the DH44 peptide and pC1 neurons exist in mammals[24, 28, 35].

## Materials and Methods

### Fly strains

Flies were raised in a 22°C or 25°C environment with 60% humidity and a 12hr:12hr light: dark cycle. Each cross used 20 virgin females with half males and was flipped every three days. In this study, all the tested flies were group-housed virgins (∼20 single-sex flies per vial) between the ages of 5-7 days after eclosion without specifically mentioned and transferred to fresh food the day before the experiments. Males and females were operated similarly. *Canton-S* and *w^1118^* flies were used as wild-type strains. *UAS-FRT-stop-FRT-CsChrimson* and *UAS-FRT-stop-FRT-GCaMP6m* were from Dr. Fang Guo; *DH44^GAL4^* and *DH44^LexA^* were from Dr. Yi Rao [60]. *DH44^AD^* was generated in this study. The following lines were from Bloomington Drosophila Stock Center (BDSC): *UAS-FRT-stop-FRT-Kir2.1* (#67686), *8xLexAop-FLP* (#55819), *UAS-myrGFP* (#32198), *UAS-dTrpA1* (#26263), *UAS-Kir2.1* (#6596), *R71G01-LexA* (#54733), *R15A01-LexA* (#54426), *R15A01-AD* (#68837), *R26E01-LexA* (#54617), *R26E01-AD* (#75740), *pC1-SS2* (#86836) and *vpoDN-SS1* (#86868). RNAi lines were from Tsinghua Fly Center (THFC) at the Tsinghua University. *UAS-FRT-stop-FRT-dTrpA1* [34], *UAS-FRT-stop-FRT-TNT^in^*, *UAS-FRT-stop-FRT-TNT* [61], *dsx^GAL4^*, *dsx^LexA^* and *dsx^DBD^*[20] were used as previously. The full genotypes of tested flies are described in Table S1.

### Generation of *DH44^AD^*

To generate the *DH44^AD^* knock-in line, the coding sequence of *p65AD* was amplified from Addgene plasmid #26234 and cloned into the attB-GMR-miniwhite vector together with the DNA sequence that had been removed from *DH44^attP^* flies as previously described[60]. The modified vector was integrated at the attP site of *DH44^attP^* flies. Successful transformant flies were selected by the 3xP3-DsRed marker and further validated by PCR. The selection marker was excised by the Cre/loxP system.

### Generation of anti-DH44 antibody

The rabbit polyclonal antibodies against the *Drosophila* DH44 protein were generated by Wuhan Abclonal Biotechnology. At least two specific pathogen-free Japanese White Rabbits were immunized with chemosynthetic polypeptides (DDGDNEGEDSYNDVGTEGVG, 281–300 amino acids of DH44). After a successful antiserum detection by ELISA, the rabbits were sacrificed, and their serum was affinity purified using the same antigen to obtain the final antibodies. The specificity of the custom anti-DH44 antibodies was verified by immunostaining of wild-type Canton-S flies. Six cells in the PI region of the brain were observed, which have been previously described[36].

### Immunostaining

The brains of 7-day-old flies were dissected in Schneider’s insect medium (Thermo Fisher Scientific, 21720). They were then fixed in 4% paraformaldehyde in phosphate-buffered saline (PBS) for 1h on ice. After being washed 4 times for 15 minutes each in PAT3, which consisted of 0.5% (v/v) Triton X-100 and 0.5% (w/v) bovine serum albumin in PBS, the brains were blocked in 3% normal goat serum (NGS) for 1h at room temperature. Brains were subsequently incubated with the primary antibody solution which had been diluted in 3% NGS overnight at 4°C, following a 4-times 15-min wash in PAT3 at room temperature. The primary antibodies used here were rabbit anti-GFP paired with nc82, which were diluted at a ratio of 1:1000 and 1:100 or rabbit anti-DH44 paired with mouse anti-GFP, which were diluted at a ratio of 1:500 and 1:1000. Finally, the brains were incubated with the secondary antibody solution using the same preparation way for 1-2 days at 4°C. All secondary antibodies used here (donkey anti-rabbit or mouse IgG conjugated to Alexa 488 and donkey anti-rabbit or mouse IgG conjugated to Alexa 555) were diluted at a ratio of 1:500. After being washed 4 times thoroughly in PAT3 at room temperature, the brains were mounted by the VECTASHIELD mounting medium for imaging. Whole brains were imaged at 10× and cell bodies were imaged at 20× lens (Olympus) using confocal microscope (Zeiss, LSM900) with a resolution of 1024 x 1024 pixels. All imaging data were analyzed using Fiji.

### *ex vivo* calcium imaging

For temperature-activated *ex vivo* calcium imaging, transgenic flies expressed both *GCaMP6m* and *dTrpA1* in pC1^R71G01^ neurons were collected after eclosion and raised for 7 days at 22°C. Male and female flies were dissected in *Drosophila* Adult Hemolymph-liked Saline (2.0 mM CaCl_2_, 5 mM KCl, 5 mM HEPES, 8.2 mM MgCl_2_, 108 mM NaCl, 4 mM NaHCO_3_, 1 mM NaH_2_PO_4_, 10 mM sucrose, 5 mM Trehalose).

To activate dTrpA1, a custom-designed heating unit was fixed on the object stage[62]. The dissected brains were placed between the heating plate and a siliconized cover glass (Diamond, 22 X 22mm) surrounded by 30μl AHL solution. A circuit board (Accuthermo, FTC200) was utilized to regulate the temperature, which was connected to a thermocouple responsible for monitoring the real-time temperature of the samples, and a semiconductor refrigeration board (CL-C067) that adjusted the temperature. The stimulation paradigm consisted of a 1-min baseline at 22°C and four consecutive 2.5-min heating events. Each event took 30 seconds to heat up and was maintained at that temperature for 2 minutes.

Fluorescence was captured by a 20× water immersion lens (Olympus) performed with a two-photon microscope (ZEISS). The focal plane with the highest number of cell bodies for each brain was scanned at a speed of 1 Hz. All imaging data were analyzed using Fiji and cell bodies were manually circled as regions of interest (ROI). ‘Mean ΔF/F’ was calculated as (F-F_0_)/F_0_, where F_0_ was the difference in average fluorescent intensity between target cell bodies and another non-target same-size region at 22°C, and F was the difference in average fluorescent intensity between target cell bodies and another non-target same-size region at each activation temperature. The average value of the ‘Mean ΔF/F’ at each temperature minus the one at 22°C was the ‘Change in ΔF/F’.

For the results shown in Figure S2, to cover the majority of female pC1 neurons, sample was scanned at a speed of 0.2 Hz while images for multiple focal planes were collected. Each cell body of Z stacked images was manually circled as ROI. Basal calcium level of pC1 cells was the average fluorescent intensity of the baseline.

### Behavioral assays

We assayed sleep, feeding, sex, aggression and spontaneous locomotion for both males and females. Flies were raised and tested at 25°C, except for the dTrpA1-related experiments, where flies were raised and kept at 22°C until tested at specific activation temperatures. Behavioral assays were started at ZT6 on the test day and digital videos were recorded by HD cameras (Sony, FDR-AX45) at 25 frames per second under a visible light if not otherwise specified. Details for each behavioral assay are listed below.

### Male courtship and female receptivity assay

Two-layer chambers with a diameter of 1cm (48-well) were used to test both male courtship and female receptivity. These two behaviors have similar ways of detection, with a difference that tested males were paired with 4 to 6-day-old wild-type virgin females in the male courtship assay while tested females were paired with 4 to 6-day-old wild-type virgin males in the female receptivity assay. On the test day, a tester and its single target were loaded individually into the chambers with gentle aspiration through a hole in the top plate. They were separated by a plastic transparent thin film in the middle of two acrylic plates until the videotaping began. In the activation experiments, loaded flies were allowed to recover for 1 hour at 22°C. Then they were pre-warmed at a specific activation temperature (27°C, 28.5°C, 30°C or 32°C) or a control temperature (22°C) for 30min with the existence of the transparent film. Once the videotaping began, two flies were encountered on account of removing the transparent barrier and recorded for another 30min. For experiments that do not involve temperature control, loaded flies were recorded right after a 1-hour recovery at 25°C. Such recording also persisted for 30min at 25°C after removing the film.

Male courtship was evaluated by two parameters including ‘courtship index’ and ‘cumulative copulation rate’. The courtship index was calculated as the fraction of time the flies spent performing any step of courtship within 10 min after courting initiation. The cumulative copulation rate was calculated as the cumulative proportion of males who copulated successfully per 5 minutes throughout the 30-min movies. For female receptivity, the cumulative percentage of copulation was counted per 2 minutes throughout the 30-min movies, and the percentage at 5min and 30min were redrawn as a histogram for further comparison. Both males and females were scored manually with the usage of a LifeSongX software.

### Wing extension assay

Single male wing extension assay was performed in activation experiments. One-layer chambers with a diameter of 2cm (12-well) were used to test such behavior. Individual males were gently aspirated into the chamber and allowed for a 1-hour recovery at 22°C. After pre-warming at experimental temperature for 1h, flies were recorded for 10min by a camera (HIK VISION). Some measures were applied to make the flies’ movements clearer, such as using an infrared backlight (Sea-light, GB200X300-RGBIR) and magnifying the images as much as possible. The side walls of the cylindrical arena were additionally coated with Insect-a-Slip (BioQuip Products, 2871B) the day before test to prevent flies from climbing up such surfaces. Wing extension was analyzed automatically (see fly tracking and behavior classification) with an evaluation indicator of wing extension index, which was calculated as the fraction of time the flies spent performing the unilateral wing extension within the indicated period.

### Aggression assay

Two-layer chambers with a diameter of 1.5cm (16-well) were used to test aggression for both sexes. Insect-a-Slip was used as above mentioned. Two tested flies of the same genotype and sex were gently aspirated into the chamber and separated by a transparent thin film. For the activation experiments, flies were group-housed until test. We used pairs of flies from separate vials so they were new to each other. Loaded flies were pre-warmed at experimental temperature for 30min following videotaping for another 30min after a 1-hour recovery at 22°C. For loss-of-function experiments, both male and female flies were individually housed in a 2ml food tube after eclosion to raise the aggressiveness naturally. In addition, a freshly prepared food patch (diameter: 8 mm; depth: 3 mm) was added in the center of each fly pair as a stimulus during the assay. The bottom food substrate consisted of regular food mixed with commercial apple juice (60% v/w) and yeast (1% w/w). Loaded flies were allowed to recover for 1 hour at 25°C and videotaped for 30min. The barrier was not removed until the camera started recording. The number of lunges for males and head-butts for females within 10 min are counted manually, in which 10 minutes start after the first initial lunge or head-butt. For males, there was an additional analysis of the percentage of tested pairs that performed high-intensity fights (holding, boxing, and tussling) within the same 10 minutes in each experiment.

### Spontaneous locomotion assay

Two-layer chambers with a diameter of 2cm (24-well) were used to test spontaneous locomotion for both sexes. Individual flies were gently aspirated into the upper layer of the chambers with light color food filled into the lower layer. In all experiments, loaded flies were placed directly into an incubator pre-set with the specific temperature at ZT0 of the test day, and a 24-hour movie was captured with full exposure to visible light. A thin film was removed upon videotaping in all experiments to make the initial locomotion more comparable. The walking of flies was tracked by the ZebraLab software system (ViewPoint Life Sciences) with data output at 1min resolution. The average speed per 5 minutes was calculated to show the locomotor activity profile within 24 hours and the mean velocity of each fly over a 24-hour period was also calculated to facilitate comparison.

### Sleep assay

Sleep of flies was measured using the DAM2 Activity Monitor (TriKinetics). Activity counts were collected in 1-min bins. On the loading day, individual flies at the age of 2-3 days after eclosion were loaded into glass tubes (5mm diameter) containing 5% sucrose and 2% agarose (w/v) and inserted into holes in the monitor case. Loaded flies were placed in a DigiTherm CircKinetics incubator (Tritech Research) entrained to 12L: 12D. Activity recordings were analyzed from ZT0 on the second day. For thermogenetic activation experiments, the test was performed at 22°C for one day as a baseline before entering experimental temperatures following another 1-day recovery at 22°C. The temperature was altered at ZT0 and stabilized within 30 min. Sleep was analyzed based on the output files with beam cross counts using custom-designed software. Dead flies were excluded from the data analysis. Data from three test days of the same flies were averaged to create a sleep profile which was defined as the fraction of time in sleep in consecutive 30-min sections. Change in total sleep was calculated as the percentage of sleep change on the day of temperature shift relative to baseline.

### Food intake assay

Food intake was measured using food labeling with a colorimetric dye. We collected 20 freshly eclosed flies and housed them for 4–6 days in a vial with standard food. Following 24-hour starvation on 1% agar (w/v), flies were fed a blue dye by incorporating it into the food they consume with a recipe calling for 50% commercial apple juice, 2.5% sucrose, 2.5% yeast extract, 0.5% agar, and 1% blue dye. For activation involving dTrpA1, starved flies were transferred to empty vials and pre-warmed at an experimental temperature or a control temperature for 30min before feeding test. For regular experiments at 25°C, flies were transferred to the vials containing dye labeling food right after starvation. Feeding was terminated after 15 minutes by transferring flies to an empty vial and freezing the vials at −80 °C immediately. The absorbance of ingested blue dye reflecting the amount of consumed food was detected to quantify the food intake. 10 frozen flies were transferred to 1.5-ml EP tubes randomly and homogenized in 500 μl 1 X PBS buffer with 1% Triton X-100. The absorbance of supernatant after 30-minute centrifugation was detected at 630nm using a NanoDrop 2000 Spectrophotometer (Thermo Scientific).

### Fly tracking and behavior classification

Analysis of wing extension behaviors in all experiments was performed by automated behavioral classifiers developed with the JAABA (Janelia Automatic Animal Behavior Annotator)[63]. Videos were first tracked using Caltech FlyTracker[64], which can provide a variety of behavioral information about the flies, and we then trained a classifier for wing extension. Its performance against ground truth data labeled manually was validated and listed in Table S2. We manually labeled whether the flies were displaying a certain behavior in a set of randomly chosen frames through the ‘ground-truthing’ mode in JAABA using videos outside of the training datasets. The precision and recall between the ground truth scores and predicted scores were additionally calculated.

### Optogenetic assay

For optogenetic experiments, crosses were reared under standard conditions at 25°C but newly eclosed flies were transferred to retinal food which was made by adding all-trans-retinal to standard fly food, with a final concentration of 1 mM and kept in the dark until the test. A LED panel with 4 channel (Sea-light, GB200X300-RGBIR) was fixed on the stage to provide photostimulation with different wavelengths of light at various intensities which was adjusted by an externally dimmable LED driver (Sea-light, GP-245-4C). The light switch was controlled by a custom-designed PLC (Uni Electronic Tech, UN2070-0606RT). Two-layer behavioral chambers with a diameter of 2cm (24-well) were placed on the surface of the LED panel. Each chamber held a single fly, which was gently aspirated into the upper layer and transparent food (5% sucrose and 2% agarose) in the lower layer to reduce heat caused by photostimulation. Flies were recorded under an infrared backlight by a camera (HIK VISION) equipped with an infrared filter at 12 fps to allow the visualization of the flies in the dark and avoid detection of light from the photostimulation.

All optogenetic experiments in our study used constant illumination with the wavelength at 620nm. Flies were loaded and acclimated in the setup for 1 hour before the test. In the experiment shown in Figure 2, we tested the same set of flies with 9 increasing intensities of 30s photostimulation with a 2-min inter-stimulus interval after a 2-min pre-stimulation baseline. The 9 levels of photostimulation are named from 1 to 9, whose intensities correspond to 0.48, 0.59, 0.70, 0.81, 0.92, 1.04, 1.15, 1.26, 1.37 μW/mm^2^, measured by an optical power meter (Fastlaser Tech) placed on the board with transparent food. For the experiment shown in Figure 5, 1-min photostimulation at level 7 (1.15 μW/mm^2^) was given after a 5-min pre-stimulation recording followed by a 10-min post-stimulation recovery. These experiments were conducted within ZT1-ZT5. The mean velocity of each stage (pre-, during- and post-stimulation) were calculated.

### Cell picking, RNA sequencing and analysis

We initially used a *P1-splitGAL4* (*R15A01-AD; R71G01-DBD*) driving *UAS-tdTomato* for manual cell picking of P1/pC1 neurons. In brief, tdTomato^+^ cells were manually picked from dissected fly brains under a fluorescence microscope following a previous protocol [65], with a minimum of 50 tdTomato^+^ cells for each sample. The picked cells underwent three washes before being loaded into a lysis buffer to ensure sample purity. RNA was extracted by the PicoPure RNA Isolation Kit (KIT0204, Thermo Scientific) and converted to cDNA by the Ovation RNA-Seq System V2 (NuGEN, #7102-A01). The Initial quality check was performed using qPCR with the *dsx* gene as a positive control and the *repo* gene as a negative control, before proceeding to DNA fragmentation and the final adaptor ligation for sequencing. The cDNA libraries were sequenced by Illumina Hiseq 2500 platform. The sequenced raw data were first pre-processed to remove low-quality reads, adaptor sequences and amplification primers. Reads were mapped to *Drosophila* genome and the mapped reads were selected for further analysis. Gene expression was quantified using TPM (transcripts per million). This preliminary data identified *DH44* as a potential molecular marker for a subset of pC1 neurons for further studies (Figure 3A).

### Statistical analyses

Statistical analyses were performed by GraphPad Prism 9. Briefly, for comparison between two groups, Unpaired t-tests or Mann-Whitney U-tests were conducted depending on if the data come from normal distribution. For comparison between multiple groups, one-way ANOVAs or Kruskal-Wallis were performed, and post hoc tests were used to compare specific pairs within groups. P>0.05 was considered to be not significant. The sample sizes (n number) and the statistical methods for each experiment are indicated in the figure legends. See Table S1 for more details.

## Acknowledgments

We thank the Bloomington *Drosophila* Stock Center and Tsinghua Fly Center for fly stocks, and Dr. Zhiyong Liu and Chao Li (Institute of Neuroscience, Chinese Academy of Sciences, China) for the help on picking pC1 neurons. This work was supported by grants from National Key R&D Program of China (2019YFA0802400) and the National Natural Science Foundation of China (31970943 and 32371067).

## Author contributions

X.J. and Y.P. conceived the study. J.C and M.S. collected brain samples for RNA sequencing, M.M. setup the temperature control system for GCaMP imaging, and X.J. performed all other experiments and analyzed the data. X.J. and Y.P. wrote the manuscript.

## SUPPLEMENTARY INFORMATION

**Figure S1.**
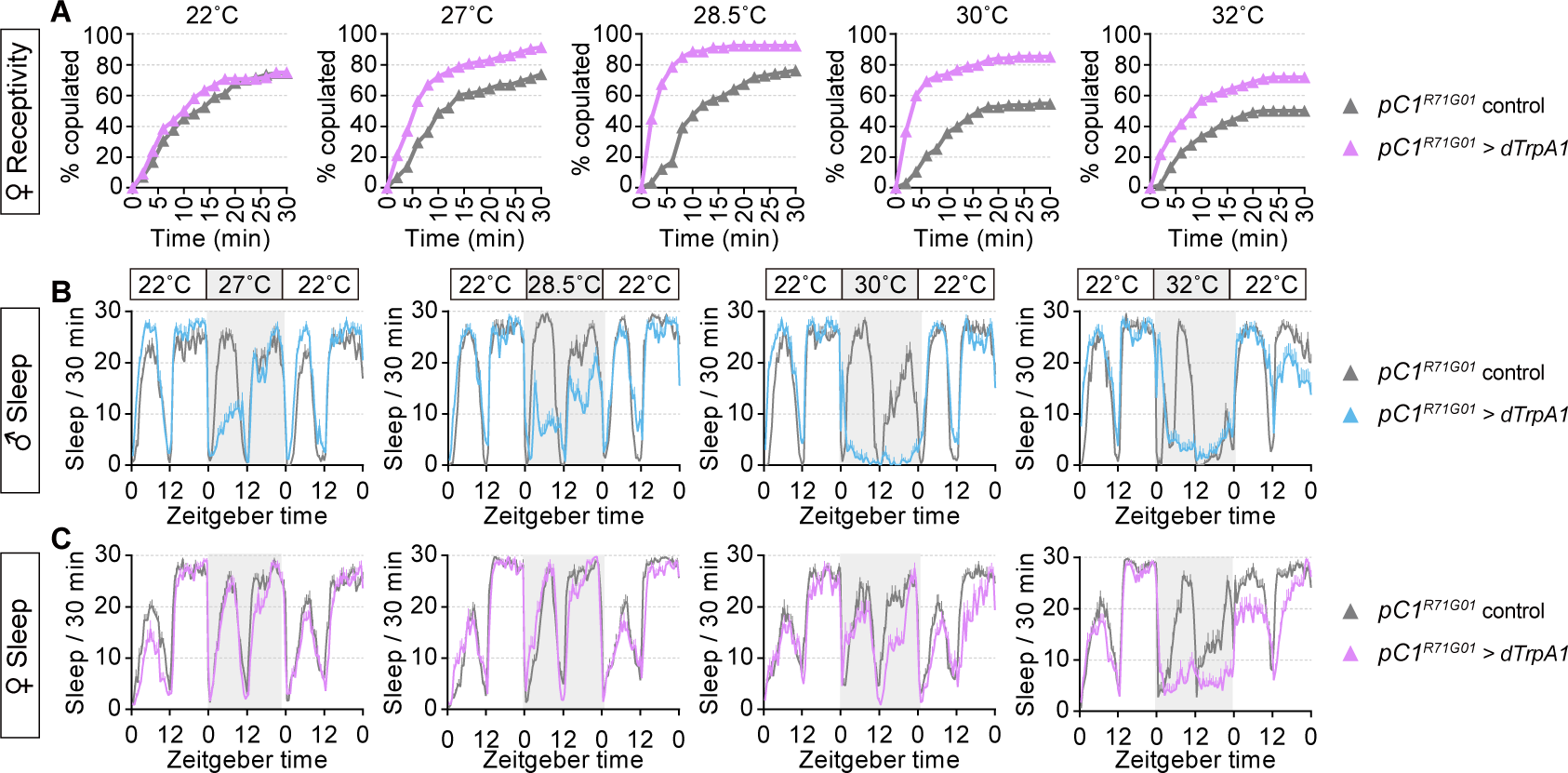
Detailed behavioral patterns of receptivity in females and sleep in both sexes after thermogenetic activation of pC1^R71G01^ neurons. (A) 30-min receptivity profile of virgin females corresponding to Figure 1D. (B and C) Sleep profile for three days in males (top) corresponding to Figure 1I, and females (bottom) corresponding to Figure 1J, respectively. See Table S1 for full genotypes of flies and sample sizes in this and all subsequent figures.

**Figure S2.**
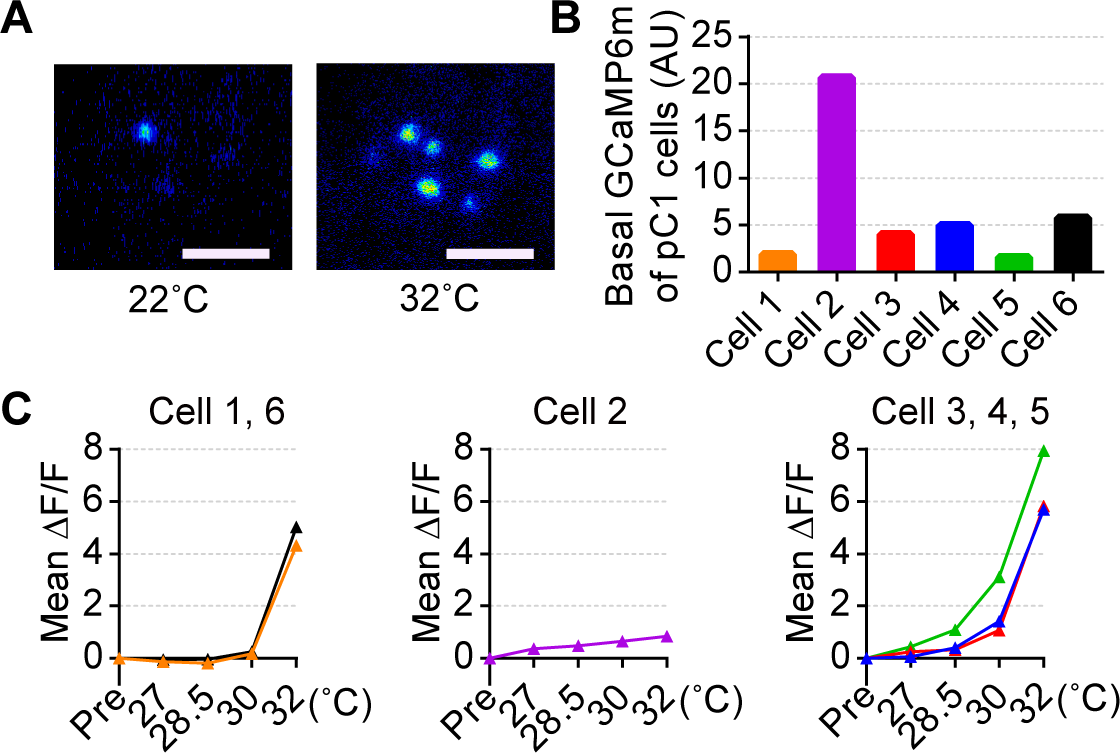
Individual female pC1 neurons have different activation properties. (A) Representative pseudo-colored images of a female brain with maximum intensity projection of fluorescence in pC1^R71G01^ neurons before and after temperature ramping. Scale bars: 30 μm. (B) Basal fluorescence intensity of GCaMP6m in individual female pC1^R71G01^ cell bodies. (C) Mean fluorescence changes (ΔF/F) in individual female pC1^R71G01^ cell bodies during each 2-min temperature maintenance stage. pC1^R71G01^ neurons were divided into three types based on their calcium responses. *ex vivo* calcium imaging experiments have a same protocol shown in Figure 1B.

**Figure S3.**
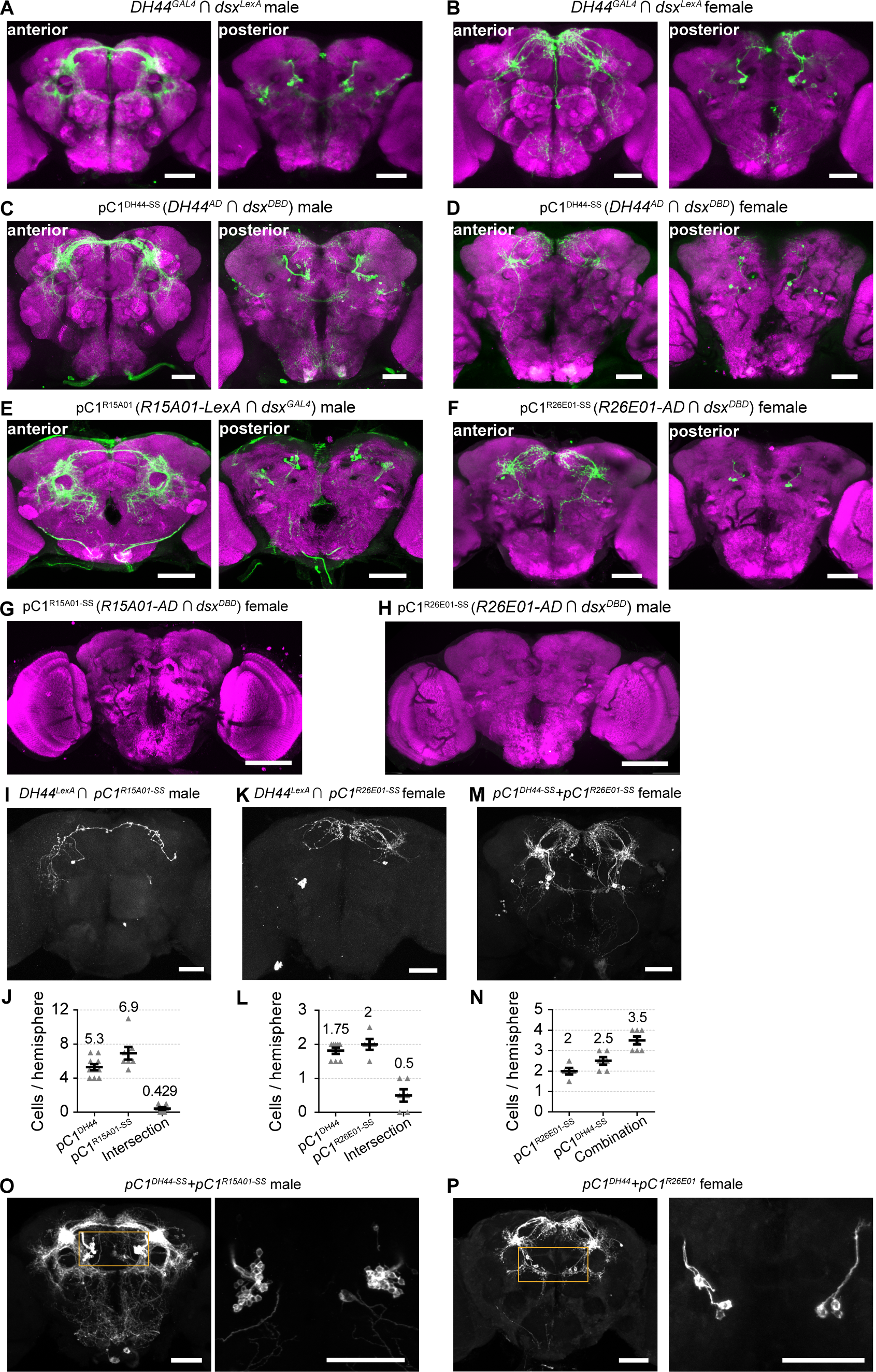
Anatomic characterization of identified pC1 subtypes. (A and B) Confocal images showing brain anterior (left) and posterior (right) parts of pC1^DH44^ neurons from a male (A), and a female (B) with the genotype of *dsx^LexA^*, *DH44^GAL4^*, *UAS-FRT-stop-FRT-CsChrimson*, and *LexAop-FLP*, co-stained with anti-GFP (green) and anti-nc82 (magenta). Scale bars: 50 μm. (C and D) Confocal images showing brain anterior (left) and posterior (right) parts of pC1^DH44-SS^ neurons from a male (C), and a female (D) with the genotype of *DH44^AD^*, *dsx^DBD^*, and *UAS-myrGFP*, co-stained with anti-GFP (green) and anti-nc82 (magenta). Scale bars: 50 μm. (E) Confocal images showing anterior (left) and posterior (right) parts of pC1^R15A01^ neurons in the *R15A01-LexA*, *dsx^GAL4^*, *UAS-FRT-stop-FRT-CsChrimson*, and *LexAop-FLP* male brain, co-stained with anti-GFP (green) and anti-nc82 (magenta). Scale bars: 50 μm. (F) Confocal images showing anterior (left) and posterior (right) parts of pC1^R26E01-SS^ neurons in the *R26E01-AD*, *dsx^DBD^*, and *UAS-myrGFP* female brain, co-stained with anti-GFP (green) and anti-nc82 (magenta). Scale bars: 50 μm. (G) Confocal image of the *R15A01-AD*, *dsx^DBD^*, and *UAS-myrGFP* female brain, co-stained with anti-GFP (green) and anti-nc82 (magenta). Scale bar: 100 μm. (H) Confocal image of the *R26E01-AD*, *dsx^DBD^*, and *UAS-myrGFP* male brain, co-stained with anti-GFP (green) and anti-nc82 (magenta). Scale bar: 100 μm. (I and J) Expression pattern (I) and cell count per hemisphere (J) of DH44-positive pC1^R15A01-SS^ neurons in *R15A01-AD*, *dsx^DBD^*, *DH44^LexA^*, *UAS-FRT-stop-FRT-CsChrimson*, and *LexAop-FLP* male brains. Scale bar: 50 μm. n=7-10 brains. (K and L) Expression pattern (K) and cell count per hemisphere (L) of DH44-positive pC1^R26E01-SS^ neurons in *R26E01-AD*, *dsx^DBD^*, *DH44^LexA^*, *UAS-FRT-stop-FRT-CsChrimson*, and *LexAop-FLP* female brains. Scale bar: 50 μm. n=5-8 brains. (M and N) Expression pattern (M) and cell count per hemisphere (N) of combinational labeling between pC1^DH44-SS^ and pC1^R26E01-SS^ neurons in *R26E01-AD*, *dsx^DBD^*, *DH44^AD^*, and *UAS-myrGFP* female brains. Scale bar: 50 μm. n=5-7 brains. (O and P) Expression pattern of combinational labeling between pC1^DH44-SS^ and pC1^R15A01-SS^ neurons in the *R15A01-AD*, *dsx^DBD^*, *DH44^AD^*, and *UAS-myrGFP* male brain (O), and pC1^DH44^ and pC1^R26E01^ neurons in the *DH44^LexA^, R26E01-LexA*, *dsx^GAL4^*, *UAS-FRT-stop-FRT-CsChrimson*, and *LexAop-FLP* female brain (P). Cell bodies in the region within a yellow square is magnified in right. Scale bars: 50 μm.

**Figure S4.**
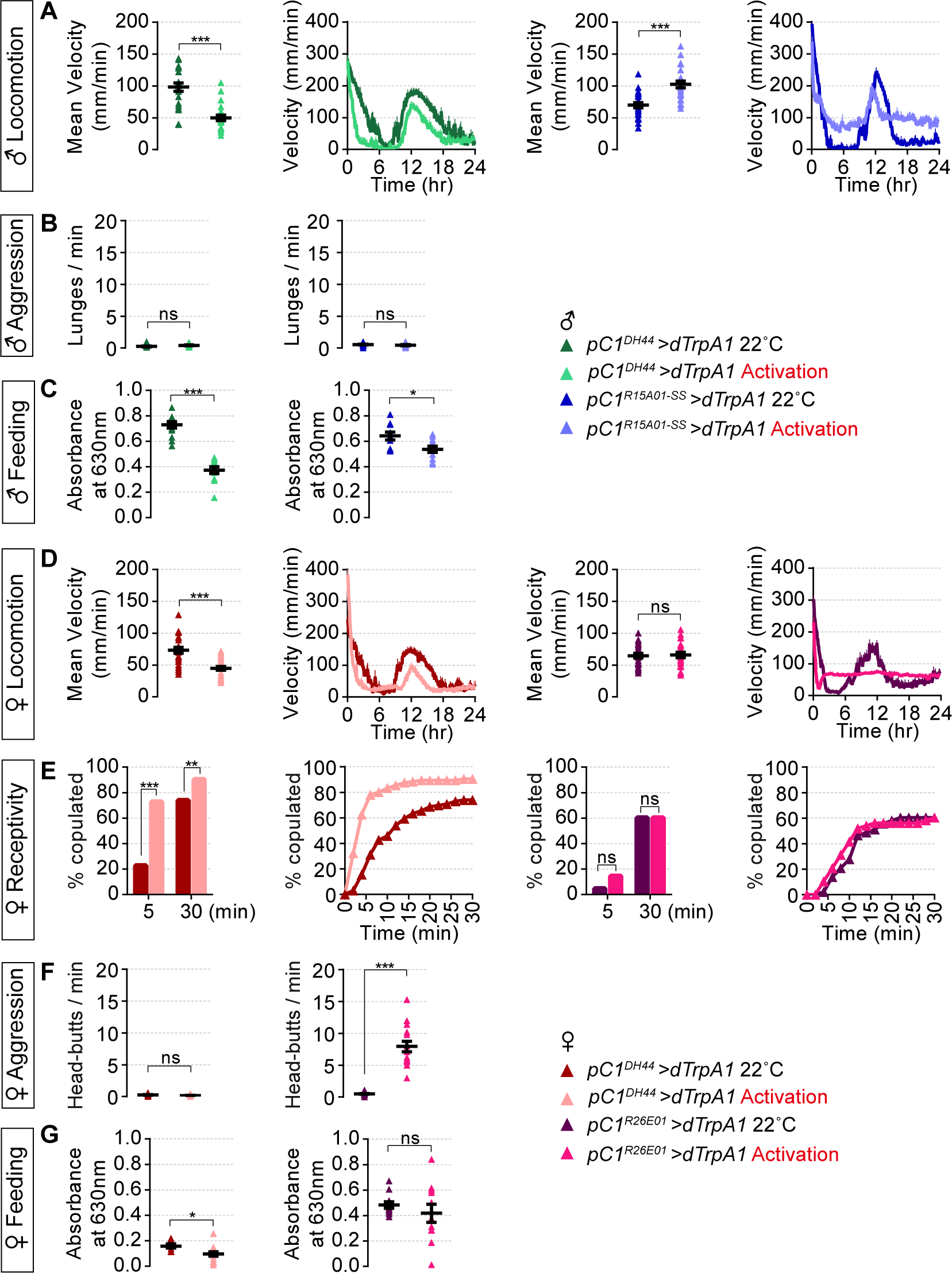
Control experiments for behavioral performance after manipulating the activity of pC1 subtypes. (A) 24-hour locomotion profile and mean velocity for *pC1^DH44^>dTrpA1* and *pC1^R15A01-SS^>dTrpA1* males at 22°C and 30°C. n=22-24. (B) Frequency of lunges for *pC1^DH44^>dTrpA1* and *pC1^R15A01-SS^>dTrpA1* males at 22°C and 30°C. n=16 pairs. (C) Food intake characterized by absorbance at 630nm for *pC1^DH44^>dTrpA1* and *pC1^R15A01-SS^>dTrpA1* males at 22°C and 30°C. n=10-13 tests, and 10 flies were used for each test. (D) 24-hour locomotion profile and mean velocity for *pC1^DH44^>dTrpA1* and *pC1^R26E01^>dTrpA1* females at 22°C and 32°C. n=23-24. (E) Receptivity curve and copulation percentage within 5 min and 30 min for *pC1^DH44^>dTrpA1* and *pC1^R26E01^>dTrpA1* females at 22°C and 27°C. n=43-96. (F) Frequency of head-butts for *pC1^DH44^>dTrpA1* and *pC1 ^R26E01^>dTrpA1* females at 22°C and 32°C. n=15-16 pairs. (G) Food intake characterized by absorbance at 630nm for *pC1^DH44^>dTrpA1* and *pC1 ^R26E01^>dTrpA1* females at 22°C and 28.5°C. n=8-18 tests, and 10 flies were used for each test. Data of activation experiments were from Figure 3 and Figure 4 for comparison with additional controls. Detailed statistical comparisons are listed in Table S1. ns, not significant, ^∗^p<0.05, ^∗∗^p<0.01, ^∗∗∗^p<0.001, error bars indicate SEM.

**Figure S5.**
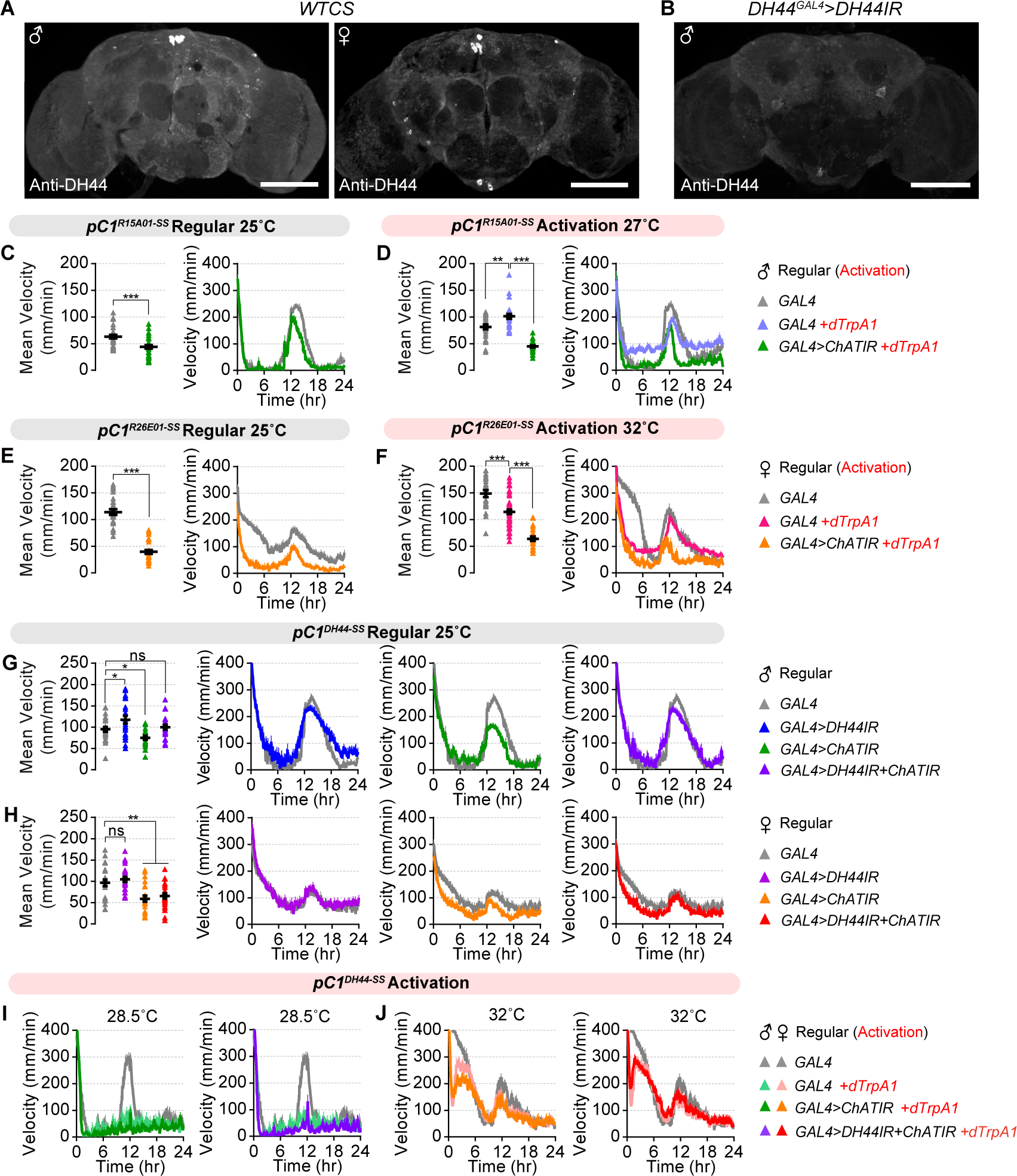
DH44 and ACh differently regulate locomotor activity in both sexes. (A) Confocal images showing DH44 immunostaining from wild-type male (left) and female (right) brains. DH44 expression is most strong in the Pars Intercerebralis (PI) region. Scale bars: 100 μm. (B) Confocal image showing DH44 immunostaining from a male fly with RNAi-mediated *DH44* knockdown under the *DH44^GAL4^* driver. Scale bar: 100 μm. (C) 24-hour locomotion profile and mean velocity in males with *ChAT* knockdown in pC1^R15A01-SS^ neurons at 25°C. n=30-32. (D) 24-hour locomotion profile and mean velocity during activation of male pC1^R15A01-SS^ neurons at 27°C with *ChAT* knockdown. n=24-29. (E) 24-hour locomotion profile and daily mean velocity in females with *ChAT* knockdown in pC1^R26E01-SS^ neurons at 25°C. n=29-31. (F) 24-hour locomotion profile and daily mean velocity during activation of female pC1^R26E01-SS^ neurons at 32°C with *ChAT* knockdown. n=21-48. (G and H) 24-hour locomotion profile and mean velocity for males (G) and females (H) with *ChAT*, *DH44*, or both knockdown in pC1^DH44-SS^ neurons at 25°C. n=20-24 for males and n=22-24 for females. (I) 24-hour locomotion profile for males during activation of pC1^DH44-SS^ neurons at 28.5°C with knockdown of *ChAT* (left), or knockdown of both *DH44* and *ChAT* (right), supplementary to Figure 5C. (J) 24-hour locomotion profile for females during activation of pC1^DH44-SS^ neurons at 32°C with knockdown of *ChAT* (left), or knockdown of both *DH44* and *ChAT* (right), supplementary to Figure 5D. Detailed statistical comparisons are listed in Table S1. ns, not significant, ^∗^p<0.05, ^∗∗^p<0.01, ^∗∗∗^p<0.001, error bars indicate SEM.

**Figure S6.**
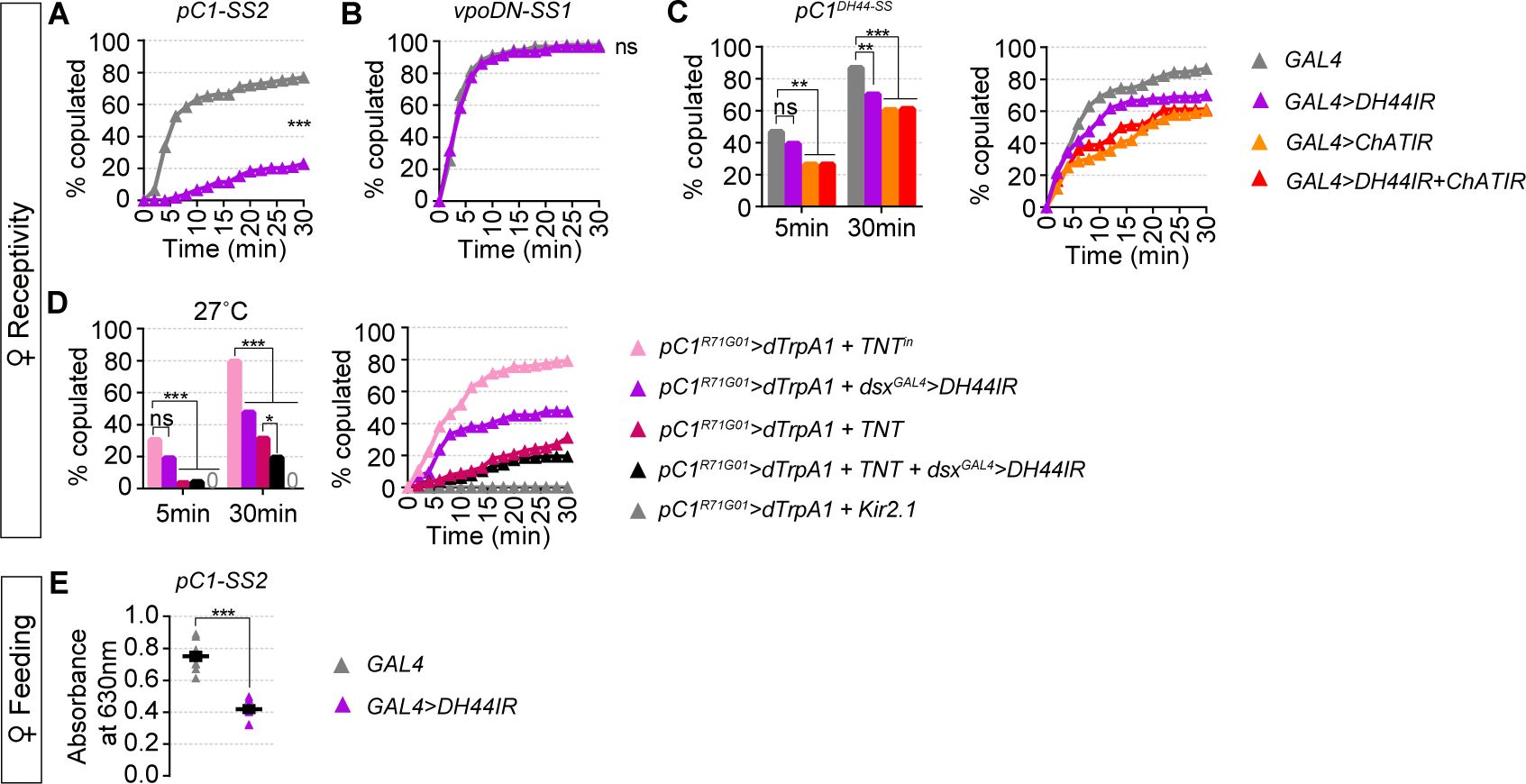
DH44 regulates female receptivity and feeding. (A and B) 30-min receptivity profile of virgin females with *DH44* knockdown in pC1-SS2 neurons (A), and vpoDN-SS1 neurons (B) at 25°C. n=93-104. (C) 30-min receptivity profile (left) and copulation percentage within 5 min and 30 min (right) for females with *ChAT*, *DH44*, or both knockdown in pC1^DH44-SS^ neurons at 25°C. n=72-90. (D) Receptivity curve and copulation percentage within 5 min and 30 min for females during activation of pC1^R71G01^ neurons at 27°C with *DH44* knockdown in *dsx^GAL4^* neurons and/or blocking synaptic neurotransmission from pC1^R71G01^ neurons by *TNT* expression. n=42-144. (E) Food intake characterized by absorbance at 630nm of females with *DH44* knockdown in pC1-SS2 neurons at 25°C. n=10-12 tests, and 10 flies were used for each test. Detailed statistical comparisons are listed in Table S1. ns, not significant, ^∗∗^p<0.01, ^∗∗∗^p<0.001, error bars indicate SEM.

Table S1. Genotypes of the flies, sample sizes, and statistical analyses. As a separate excel file.

Table S2. Behavioral classifier performance. As a separate excel file.

